# Functional characterization of gene regulatory elements and neuropsychiatric disease-associated risk loci in iPSCs and iPSC-derived neurons

**DOI:** 10.1101/2023.08.30.555359

**Authors:** Xiaoyu Yang, Ian R. Jones, Poshen B. Chen, Han Yang, Xingjie Ren, Lina Zheng, Bin Li, Yang Eric Li, Quan Sun, Jia Wen, Cooper Beaman, Xiekui Cui, Yun Li, Wei Wang, Ming Hu, Bing Ren, Yin Shen

## Abstract

Genome-wide association studies (GWAS) have identified thousands of non-coding variants that contribute to psychiatric disease risks, likely by perturbing *cis*-regulatory elements (CREs). However, our ability to interpret and explore their mechanisms of action is hampered by a lack of annotation of functional CREs (fCREs) in neural cell types. Here, through genome-scale CRISPR screens of 22,000 candidate CREs (cCREs) in human induced pluripotent stem cells (iPSCs) undergoing differentiation to excitatory neurons, we identify 2,847 and 5,540 fCREs essential for iPSC fitness and neuronal differentiation, respectively. These fCREs display dynamic epigenomic features and exhibit increased numbers and genomic spans of chromatin interactions following terminal neuronal differentiation. Furthermore, fCREs essential for neuronal differentiation show significantly greater enrichment of genetic heritability for neurodevelopmental diseases including schizophrenia (SCZ), attention deficit hyperactivity disorder (ADHD), and autism spectrum disorders (ASD) than cCREs. Using high-throughput prime editing screens we experimentally confirm 45 SCZ risk variants that act by affecting the function of fCREs. The extensive and in-depth functional annotation of cCREs in neuronal types therefore provides a crucial resource for interpreting non-coding risk variants of neuropsychiatric disorders.

## Main

*Cis*-regulatory elements (CREs) play a fundamental role in regulating cell-type-specific gene expression^1,2^. Great strides have been made in recent years in the annotation of candidate CREs (cCREs) in the human genome through the profiling of their biochemical signatures such as chromatin accessibility, histone modifications, transcription factor (TF) binding, and DNA hypomethylation, etc^3–5^, but a shortage of direct functional evidence makes it hard to know if and when the annotated cCREs indeed regulate transcription, emphasizing the urgent need to comprehensively characterize cCREs. The cCREs harbor a disproportionately large number of genetic variants associated with complex human traits and common diseases, supporting the hypothesis that non-coding variants contribute to human diseases and traits largely through the modulation of gene expression^6,7^. In particular, genome-wide association studies (GWAS) have identified thousands of non-coding genetic variants associated with neuropsychiatric disorders such as schizophrenia^8^ (SCZ), attention deficit hyperactivity disorder^9^ (ADHD), autism spectrum disorder^10^ (ASD), and bipolar disorder^11^ (BD), etc. The genetic heritability of DNA variants associated with neuropsychiatric traits is significantly enriched within cCREs annotated in the glutamatergic neurons compared to non-neuronal cell types^12–14^. However, a dearth of annotated functional CREs (fCREs) in physiologically relevant cell types such as neurons has hindered our ability to identify the disease-causing non-coding variants.

To address this knowledge gap, we utilized human induced pluripotent stem cells (iPSCs) and their excitatory neuron differentiation process to perform genome-scale CRISPRi screens combined with multiomic analysis to identify both cCREs and fCREs involved in pluripotency and neuronal differentiation. Through functional characterization of 22,000 cCREs, we identified 2,847 fCREs required for iPSC fitness and 5,540 fCREs for neuronal differentiation. We further elucidated the epigenomic features of the fCREs through integrative analysis of matched transcriptome, open chromatin regions, and 3D chromatin interactions. Finally, we demonstrated that fCREs required for neuronal differentiation are strongly enriched with non-coding risk variants linked to schizophrenia, autism, and several other psychiatric disorders, and validated 110 of them through high-throughput prime editing experiments. Our study offers new insights into the regulatory mechanisms of iPSC fitness and neuronal differentiation, with significant implications for understanding the impact of neuropsychiatric disease-associated non-coding variants in disease etiology.

### Genome-scale CRISPRi screens for iPSC proliferation and neuronal differentiation

To systematically identify fCREs, we performed pooled CRISPRi screens based on iPSC fitness and excitatory neuron differentiation phenotypes. We introduced a CAG-dCas9-KRAB cassette at the *CLYBL* safe harbor locus in a transgenic iPSC line (i^3^N-WTC11 iPSCs) (**Extended Data Fig. 1a**). The i^3^N-WTC11 iPSC line contains an integrated, isogenic, and doxycycline-inducible neurogenin-2 (*Ngn2*) cassette at the *AAVS1* safe-harbor locus, enabling rapid induction of excitatory neuron differentiation in a synchronized fashion following doxycycline treatment^15^. After CAG-dCas9-KRAB knock-in, we showed that dCas9 expression level in the i^3^N-dCas9-WTC11 iPSC line is comparable to that of a known doxycycline-inducible dCas9-KRAB iPSC line^16^ (**Extended Data Fig. 1b**), and that CRISPRi exhibits effective knockdown efficiencies in both iPSCs and induced excitatory neurons (**Extended Data Fig. 1c**). Based on these results, we proceeded with genome-scale cCREs screens using this i^3^N-dCas9-WTC11 iPSC line.

For the iPSC fitness screen, we prioritized 16,670 cCREs located within 500 Kbp from the transcription start sites (TSSs) of 1,301 genes previously identified as essential for iPSC fitness^17,18^. These cCREs overlap with open chromatin regions and H3K27ac occupancy in iPSCs, determined using ATAC-seq and ChIP-seq^19^, respectively (**Fig. 1a**, **Supplementary Table 1a**). For the neuronal differentiation screens, we first selected open chromatin regions identified in iPSC-derived excitatory neurons following a 2-week differentiation, which are located within 500 Kbp of 269 genes previously identified as essential for neuronal differentiation or survival using the same differentiation system^20^. Notably, 170 of the 269 genes (63.2%) were also identified as essential for iPSC fitness in the same study, suggesting that essential genes can be shared between iPSCs and differentiated neurons^20^. Because these genes were identified from a screen of 2,325 druggable genes, which may not cover all essential genes for neuronal differentiation and fitness, we further expanded our list to include cCREs overlapping with neuronal open chromatin regions and H3K27ac sites^21^ within 500 Kbp of 1,301 iPSC essential genes. Finally, we added 282 neural enhancers annotated by mouse transgenic experiments^22^. In total, 14,289 cCREs were tested in the neuronal differentiation screens (**Fig. 1b**, **Supplementary Table 1b**). Altogether, we tested 7,126 TSSs and 14,874 distal cCREs in iPSC and neuronal screens, among which 5,000 TSS cCREs and 3,959 distal cCREs (**Methods**) are shared in both screens (**Fig. 1c**), allowing us to assess their context dependent function.

**Figure 1.**
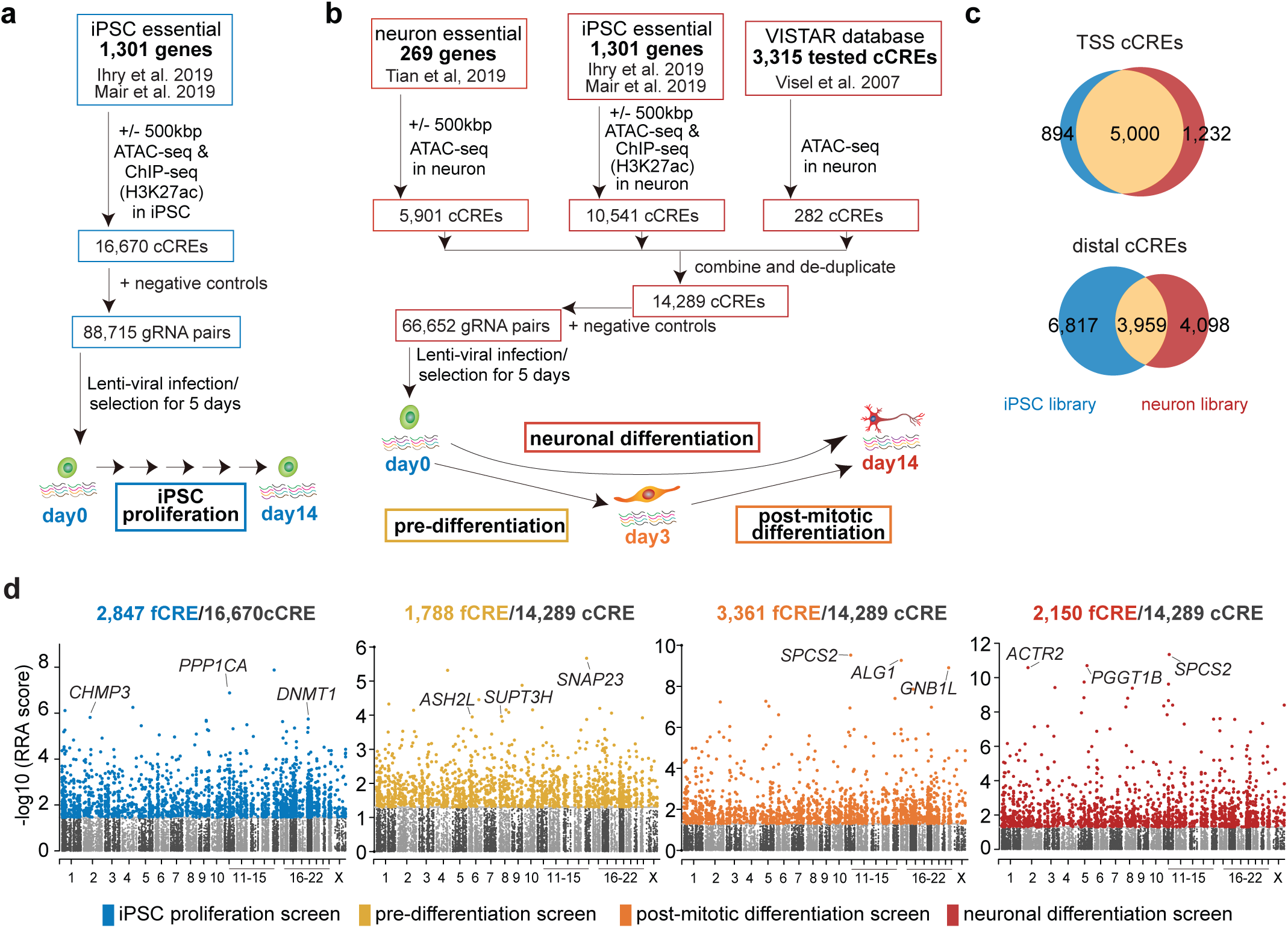
Design of CRISPRi screens to identify the fCREs. **a**, gRNA library design and screening scheme for iPSC self-renewal fCREs. **b**, gRNA library design and screening scheme for neuronal differentiation fCREs. **c**, Overlap of TSSs and distal cCREs assessed in CRISPRi screens in iPSC and neuronal cells. **d**, Genome-wide view of fCREs for iPSC renewal and neuronal differentiation. Top TSS fCREs are labeled. Positive hits are determined by FDR < 0.05.

For the CRISPRi screens, we employed the dual-gRNA CRISPRi approach as we previously demonstrated more effective dCas9-KRAB-mediated epigenetic silencing using dual-gRNAs targeting a cCRE than using a single gRNA^23^. For each cCRE we designed 5 pairs of gRNAs with an average genomic distance of ∼300 bp between each pair of gRNAs. In addition, we included 8,389 gRNA pairs targeting safe-harbor genomic regions and 1,011 non-targeting gRNA pairs as negative controls^24,25^. In total, we constructed two lentiviral libraries expressing 88,715 and 66,652 pairs of gRNAs for iPSCs and neuronal differentiation screens, respectively (**Extended Data Fig. 1d**-**e**, **Supplementary Table 1c-d**). These dual-guide gRNAs cover 1,120 essential gene loci (+/-500 Kbp for each locus), encompassing a total of ∼1 billion base pairs, or nearly one third of the human genome.

To identify fCREs required for iPSCs fitness, we infected i^3^N-dCas9-WTC11 iPSCs at a low multiplicity of infection (MOI) (MOI = 0.5) with the lentiviral library expressing dual-gRNA pairs targeting iPSC cCREs. After five days of puromycin selection (day 0), iPSCs were expanded for another two weeks (day 14). We collected cells at day 0 and 14, amplified the integrated dual-gRNA pairs, and determined copy numbers for each dual-gRNA pair at each time point by deep sequencing (**Fig. 1a**). To identify fCREs essential for neuronal differentiation, we infected i^3^N-dCas9-WTC11 iPSCs with the lentiviral library expressing dual-gRNA pairs targeting neuronal cCREs (MOI = 0.5). After five days of puromycin selection (day 0), neuronal differentiation was initiated by doxycycline-induced *Ngn2* expression. We collected cells at three time points: day 0 (iPSCs), day 3 (pre-differentiated to post-mitotic transition), and day 14 (post-mitotic excitatory neurons). To identify fCREs essential for neuronal differentiation, we compared the copy numbers of dual-gRNA pairs between day 0 and day 3 for pre-differentiation, day 3 and day 14 for post-mitotic differentiation, and day 0 and day 14 for overall neuronal differentiation. By analyzing these three pairwise time point comparisons, we delineate the temporal activity of fCREs during differentiation (**Fig. 1b**). All screens were performed with two biological replicates, with high reproducibility observed between replicates (**Extended Data Fig. 1f**-**g**).

Using the robust ranking aggregation (RRA) method from the MAGeCK pipeline^26^, we identified 2,847 fCREs as significantly depleted after 14 passages, indicating their essential role in maintaining iPSC fitness. Similarly, we identified 1,788 fCREs needed for pre-differentiation, 3,361 fCREs for post-mitotic differentiation, and 2,150 fCREs for neuronal differentiation (FDR < 0.05) (**Fig. 1d, Supplementary Table 2a-d**). As expected, promoter regions of known essential genes were among the top hits in each screen (**Fig. 1d**). Specifically, a recent publication^27^ reported 1,520 essential neuronal genes during *Ngn2-*induced excitatory neuron differentiation. Of these, 968 were included in our design (**Extended Data Fig. 1h**). We observed a strong correlation between our study’s gRNA count fold changes for these genes and those of the recent study (**Extended Data Fig. 1i**), with 686 (69.5%) also identified as essential in our study (**Extended Data Fig. 1j**). These observations provide an external validation of the efficacy of our approach for elucidating the functional sequences essential for neuronal differentiation.

As further independent evidence for the fCREs identified in the CRISPRi screens, we performed CRISPRi experiments to verify the requirements of 6 fCREs in iPSC fitness. Confirming the genome-scale CRISPRi screen results, perturbation of one distal fCRE for *SOX2* (**Fig. 2a**) and two distal fCREs for *MYC* (**Fig. 2b**) led to the downregulation of *SOX2* and *MYC* (**Fig. 2c**) and significantly reduced iPSC survival rate (**Fig. 2d**). We also identified fCREs at *GDF3*, *DUS1L* and *MRPS23* promoters, that potentially regulate *NANOG*, *FASN* and *SRSF1* expression via long-range chromatin interaction (**Extended Data Fig. 2a**-**c**). *GDF3*, *DUS1L* and *MRPS23*, while expressed in iPSC, were not iPSC essential genes based on the previous small hairpin RNA (shRNA) screen^28^ and CRISPR knockout screens^17,18^. In this stidy, CRIPSRi targeting these promoter-proximal fCREs led to down-regulation of *NANOG*, *FASN* and *SRSF1* and decreased cell survival (**Extended Data Fig. 2d**-**e**). These examples highlighted that our fCREs enable the annotation of regulatory sequences at non-essential gene promoter regions that may work as enhancers to regulate other iPSC fitness genes.

**Figure 2.**
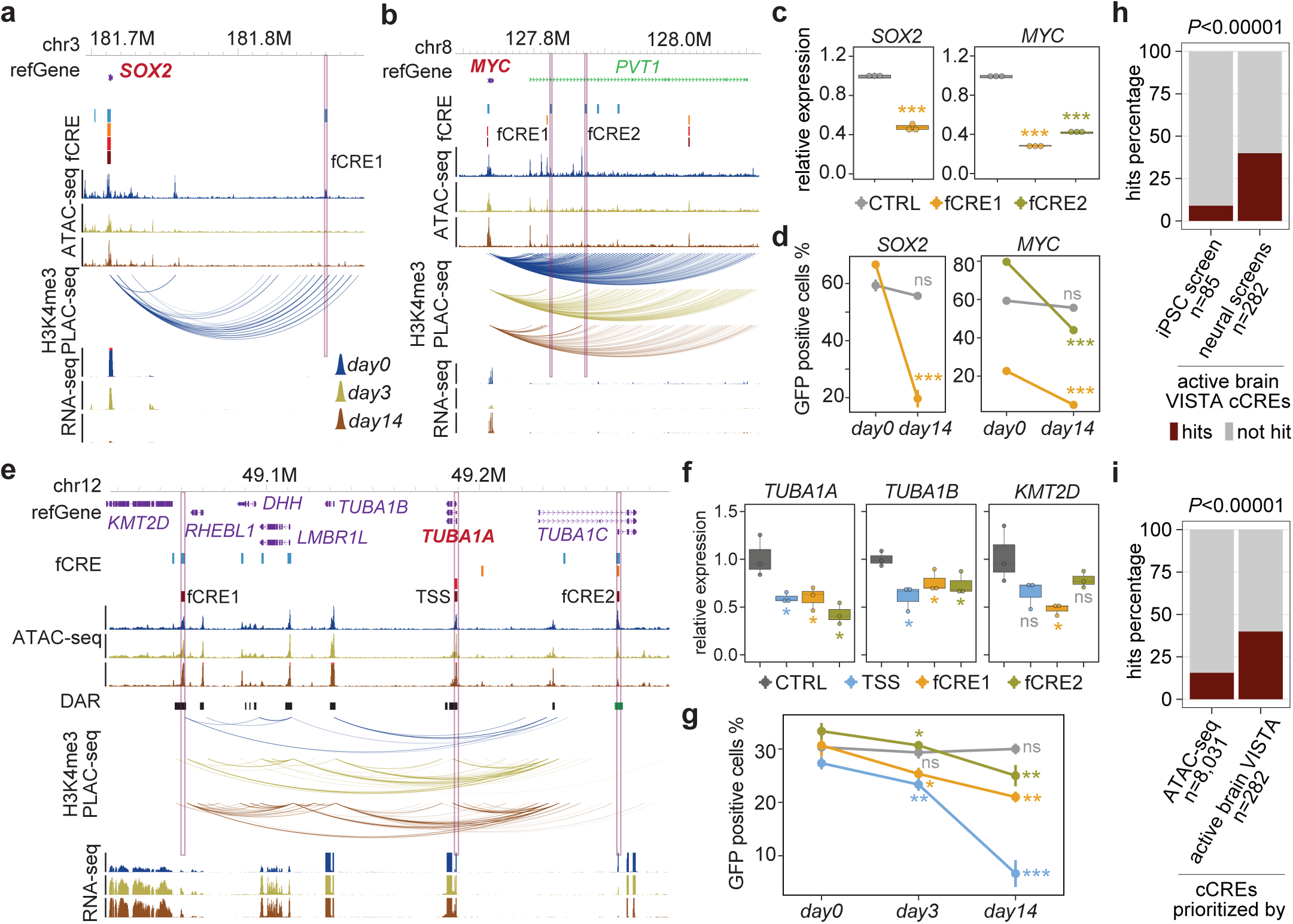
Independent verification of fCREs required for fitness of iPSC and neuronal differentiation. **a-b**, Genome browser snapshots of the *SOX2* and *MYC* loci. fCREs from the iPSC, pre-differentiation, post-mitotic, and neuronal screens are colored blue, orange, red, and dark red respectively. Purple boxes indicate the validated fCREs. **c**, CRISPRi effect of perturbing fCREs in iPSC quantified by RT-qPCRs. Boxplots indicate the median and interquartile range. Whiskers indicate the 5th and 95th percentiles. (n = 3, two-tailed two sample t-test). **d**, Survival ratio of fCRE perturbed cells by FACS in iPSC. Data are presented as mean ± s.d. (n = 3, two-tailed two sample t-test). **e**, Genome browser snapshot of the *TUBA1A* locus. fCREs from the iPSC, pre-differentiation, post-mitotic, and neuronal screens are colored blue, orange, red, and dark red respectively. Purple boxes indicate the validated fCREs. **f**, CRISPRi effect of perturbing fCREs in neurons quantified by RT-qPCRs. Boxplots indicate the median and interquartile range. Whiskers indicate the 5th and 95th percentiles. (n = 3, two-tailed two sample t-test). **g**, Survival ratio of fCRE perturbed cells by FACS in neuronal differentiation. Data are presented as mean ± s.d. (n = 3, two-tailed two sample t-test, ns: *P* > 0.05, *: *P* < 0.05, **: *P* < 0.01***: *P* < 0.001). **h**, Barplot comparing hits percentage for active brain VISTA cCREs in iPSC or neural screens. **i**, Barplot comparing hits percentage for cCREs prioritized by ATAC-seq signal or active brain VISTA cCREs in neural screens. *P* values are calculated with a two-sided Chi-squared test in (h) and (i).

We also carried out similar CRISPRi experiments to confirm the role of 4 distal fCREs identified in CRISPRi screens in the neurons. For example, the α-tubulin gene, *TUBA1A*, plays a crucial role in neural development^29^ and *TUBA1B* (Tubulin Alpha 1b) is an essential gene for neuronal differentiation^27^. Perturbing fCRE1 near the promoter of *KMT2D* or fCRE2 overlapping an alternative promoter of *TUBA1C* (**Fig. 2e**) led to the downregulation of *TUBA1A*, *TUBA1B*, and *KMT2D* (**Fig. 2f**) and affected neuronal differentiation (**Fig. 2f**). In another example, *EPHB1* encodes a protein belonging to the Ephrin-B family, which is indispensable for neurogenesis^30^. Two fCREs identified in the intronic region of *EPHB1* (**Extended Data Fig. 2f**) gain chromatin interaction with the *EPHB1* promoter in neurons compared to in iPSCs. Perturbation of *EPHB1* TSS, fCRE1, and fCRE2 affected neuronal differentiation and led to downregulation of *EPHB1* expression (**Extended Data Fig. 2g-h**).

As a final piece of evidence supporting the function of the fCREs identified in our CRISPRi screens, we compared the fCREs with previously reported tissue-specific enhancers discovered using transgenic mice^22^. These experimentally validated fetal brain enhancers (n = 282) are significantly overrepresented in the fCREs found in our neuronal CRISPRi screens (113 out of 282) (*P* < 0.01×10^-3^) but not the iPSC screen (8 out of 85) (**Fig. 2h**). Furthermore, fCREs are more enriched for the neural enhancers identified in transgenic mice (113 out of 282) than the cCREs predicted using ATAC-seq data (1,285 out of 8,031) (*P* < 0. 01×10^-3^) (**Fig. 2i**), supporting the value of fCREs.

### fCREs are strongly enriched for active chromatin marks

Our screens identified many novel fCREs in iPSCs fitness (n = 2,516) and neuronal differentiation (n = 4,887), respectively. Specifically, we identified novel fCREs at both TSSs (iPSC: n = 1,264; neuronal screens: n = 2,011) and distal elements (iPSC: n = 1,252; neuronal screens: n = 2,876) (**Fig. 3a, Extended Data Fig. 3a**). Newly identified TSS-proximal fCREs may play a role in cell fitness by directly influencing transcription from these TSSs, or by regulating other nearby genes *in cis*. Indeed 7.5% (n = 121) and 3.5% (n = 98) of the identified TSS fCREs have low expression (RPKM < 1) in iPSC and neuronal differentiation time points respectively, suggesting potential functional roles for those fCREs to act as regulatory sequences for other genes (**Supplementary Table 2e**). Strikingly, compared to TSSs of known essential genes, these novel functional TSS proximal and distal fCREs exhibit significantly lower RRA scores for postmitotic and neuronal differentiation processes (**Extended Data Fig. 3b**), demonstrating high sensitivity of dual-gRNA CRISPRi screens in detecting both TSS and distal fCREs, with perturbation of known essential gene TSSs exhibiting larger effect sizes compared to other newly identified fCREs.

**Figure 3.**
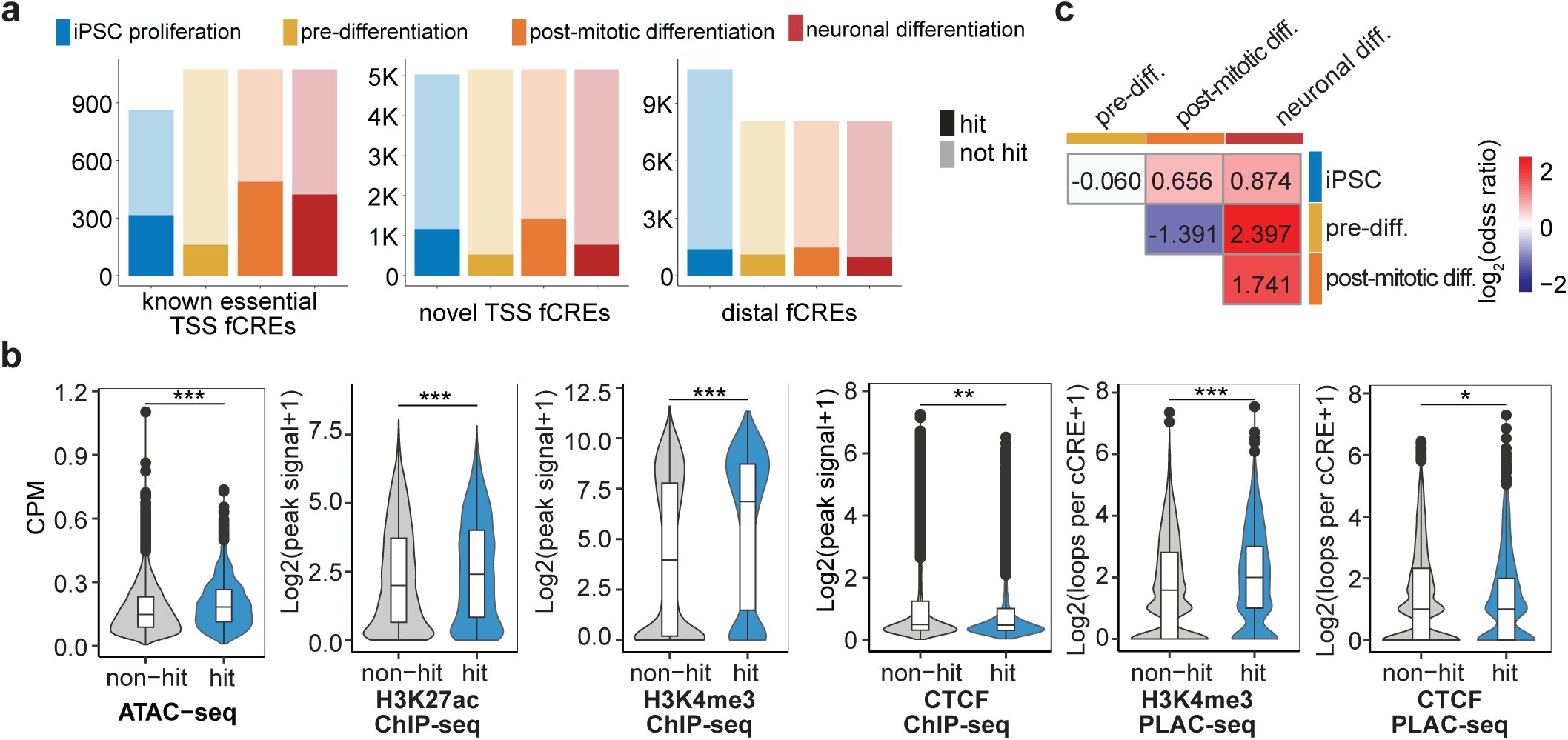
Genomic and epigenomic features of fCREs. **a**, Total amount of fCREs recovered from the previously known essential TSSs, novel TSSs, and distal fCREs for each screen. **b**, Violin plots of epigenomic signals between non-fCREs (n = 13,823) and fCREs (n = 2,847) from the iPSC screen. (Wilcoxon test, *: *P* < 0.05, **: *P* < 0.01, ***: *P* < 0.001). **c**, Heatmap of odds ratio showing the association between screens. Positive associations (OR > 1, red) are observed between the neuronal screen and either pre-differentiation or post-mitotic differentiation screen. Negative associations (OR < 1, blue) occur between pre-differentiation and post-mitotic screens.

fCREs are associated with active chromatin marks including chromatin accessibility, H3K27ac and H3K4me3 in iPSCs fCREs, evidenced by a greater enrichment of active chromatin states compared to other cCREs (**Fig. 3b)**. Moreover, iPSCs fCREs participated in on average 27.3% more H3K4me3-associated interactions per cCRE, evidenced by Proximity-ligation Assisted ChIP followed by sequencing (PLAC-seq, aka HiChIP) experiments. By contrast, CTCF signal at fCREs decreases by 10.6% despite the number of average CTCF-associated interactions at fCREs slightly increasing by 2.5% (**Fig. 3b)**.

### Neuronal fCREs display temporal changes in 3D chromatin organization during differentiation

By dividing neuronal screens into pre- and post-mitotic differentiation stages, we characterized the temporal dynamics of fCREs during neuronal differentiation. The diversity of fCREs detected throughout neuronal differentiation are highlighted by the low overlap in the fCREs found in the pre-differentiation and post-mitotic screens, with an odds ratio of a cCRE being a fCRE in both screens of 0.38 (*P* < 2.2E-16) (**Fig. 3c, Extended Data Fig. 3c**). To investigate the importance of temporal changes in the 3D epigenome for fCREs identified throughout neuronal differentiation, we conducted RNA-seq, ATAC-seq, and PLAC-seq using antibodies against H3K4me3 and CTCF at differentiation time points corresponding to our screens (**Extended Data Fig. 4a-d**, **Supplementary Table 3**). Interestingly, we observed an increase in both numbers and genomic distances of chromatin interactions after differentiation (**Fig. 4a-b**). This finding is consistent with a previous study of dynamic chromatin organization during mESC neural differentiation^31^, suggesting that terminally differentiated neurons could rely on distal gene regulation compared to undifferentiated iPSCs. Additionally, fCREs are significantly associated with chromatin interaction associated with both H3K4me3 and CTCF in the iPSC (Fisher’s exact test, *P* < 0.01), post-mitotic differentiation (Fisher’s exact test, *P* < 0.001), and neuronal differentiation screens (Fisher’s exact test, *P* < 0.05), but are negatively associated in the pre-differentiation screen (Fisher’s exact test, *P* < 0.001) (**Fig. 4c**, **Extended Data Fig. 4e**, **Supplementary Table 4a-b**). fCREs also participate in more and longer-range chromatin interactions during the differentiation process (**Fig. 4d-f**). For the 17.8% and 24.3% of fCREs that don’t engage in H3K4me3- or CTCF-associated chromatin interactions, they tend to reside further away (two-sample t-test, two-tailed, *P* < 0.001) from the nearest TSS than those distal fCREs constantly participating in H3K4me3- or CTCF-associated chromatin interactions throughout differentiation (**Fig. 4g-h)**.

**Figure 4.**
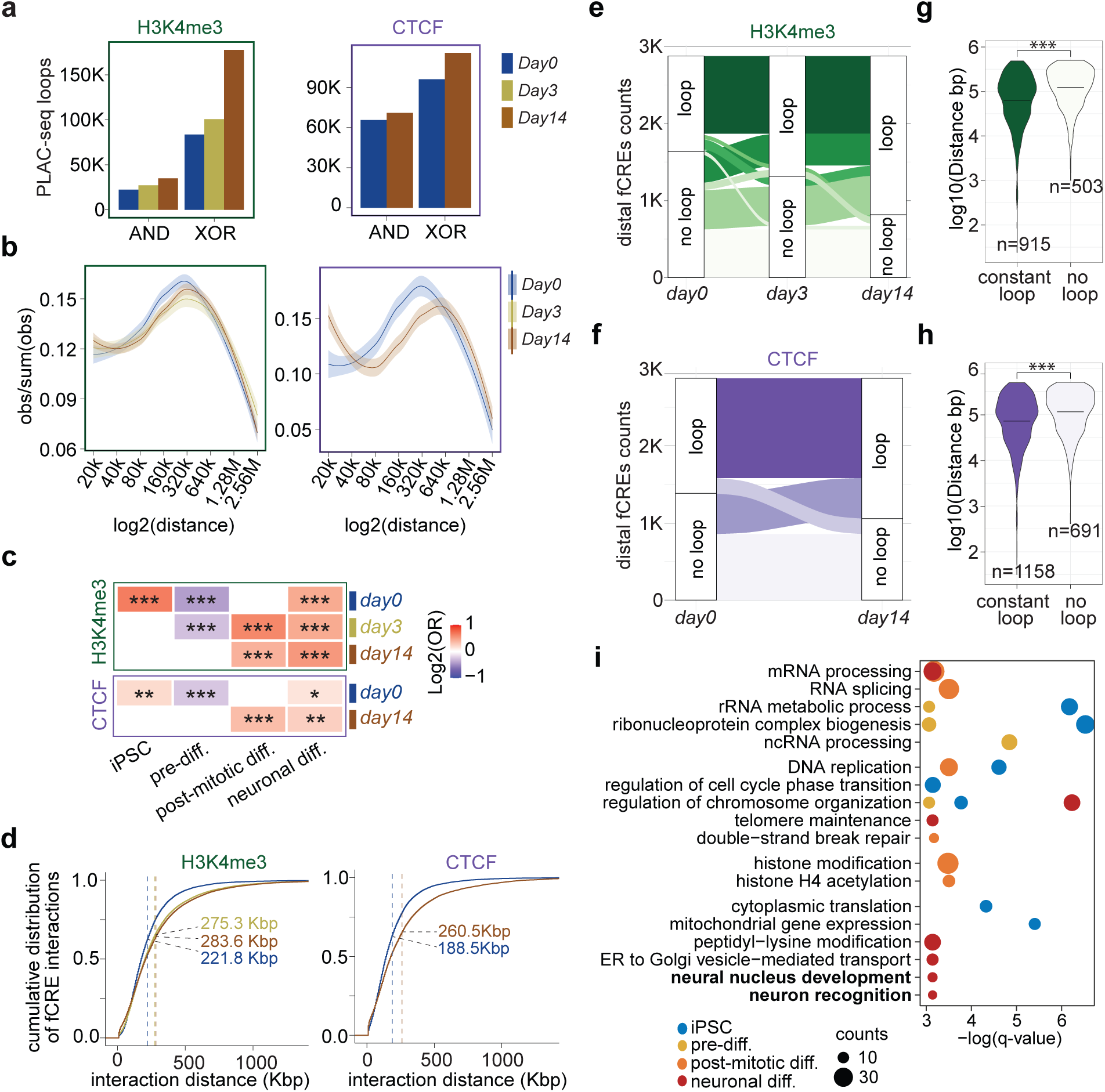
Dynamic 3D chromatin contacts at fCREs during neuronal differentiation. **a**, Chromatin contacts of H3K4me3- and CTCF-PLAC-seq experiments at each time point along neuronal differentiation. AND interactions are both the interaction bins overlap with anchor bins. XOR interactions are only one of the interaction bins overlap with anchor bins. **b**, Distance distribution of H3K4me3- or CTCF-associated contacts revealing increased 1D distances through differentiation. **c**, Heatmap showing the association between fCREs and chromatin contacts (N are indicated in **Supplementary Table 4a**, Fisher’s exact test, *: *P* < 0.05, **: *P* < 0.01, ***: *P* < 0.001). **d**, Cumulative distribution function (CDF) plots showing the median interaction distance (dash lines) shift increase through differentiation in H3K4me3- or CTCF-associated chromatin contacts containing fCREs. **e-f**, Numbers of distal fCREs that are involved in H3K4me3- (e) or CTCF-(f) associated chromatin interactions through differentiation. **g-h**, Distance of distal fCREs to their nearest TSS fCRE between fCREs in constant H3K4me3-(g) or CTCF-(h) associated interactions compared to fCREs not participating interactions (midline indicates the median, two-tailed two sample t-test, ***: *P* < 0.001). **i**, Pathway enrichment analysis for fCRE target genes (iPSC proliferation, n = 385; pre-differentiation, n = 229; post-mitotic differentiation, n = 715; neuronal differentiation, n = 438) annotated by H3K4me3 associated 3D chromatin interactions (q-value < 0.05).

To elucidate the potential biological roles of fCREs required for neuronal differentiation, we conducted gene ontology (GO) analyses. Both known and newly identified essential genes in iPSC and neuronal screens exhibit an enrichment in fundamental cellular function terms, including mRNA processing, DNA metabolic process, chromatin remodeling and mitochondrial functions (**Extended Data Fig. 4f-g**, **Supplementary Table 5a-b**). Notably, target genes of neuronal distal fCREs, evidenced by H3K4me3-centered chromatin contacts, are enriched not only in basic cellular functions, but also in such processes as neural nucleus development and neuronal cell recognition (**Fig. 4i, Supplementary Table 5c-d**). Similarly, target genes predicated by CTCF-associated interactions were enriched in processes including synapse organization, neuronal cell recognition, axon guidance, and cell-cell adhesion (**Extended Data Fig. 4h**, **Supplementary Table 5e-f**). Interestingly, these biological processes were not enriched in potential target genes predicted using distance-based annotations by GREAT^32^ (**Extended Data Fig. 4i**, **Supplementary Table 5g**). We believe that targeting gene annotation by chromatin interaction could better illuminate the role of fCREs in regulating the diverse range of biological pathways during neuronal differentiation.

To characterize the dynamic chromatin organization at fCREs during neuronal differentiation, we analyzed the PLAC-seq data and determined 9,445, 13,256, and 18,644 differentially interacting regions (DIRs) centered at promoters marked by constitutive H3K4me3 signals for pre-differentiation (day 0 vs. day 3), post-mitotic (day 3 vs. day 14), and neuronal differentiation (day 0 vs. day 14), respectively. We also determined 7,962 DIRs centered at CTCF binding sites for neuronal differentiation (day 0 vs. day 14) (**Extended Data Fig. 5a**, **Supplementary Table 6**, details in **Methods**). Overall, DIRs are positively correlated with transcriptional changes at their target promoters and distal accessible regions connecting to these promoters display changes in accessibility congruent with transcription from the promoters (**Fig. 5a, Extended Data Fig. 5b**). Furthermore, DIRs centered at promoters are enriched for fCREs identified in the post-mitotic and neuronal differentiation screens but depleted in pre-differentiation fCREs, compared to the rest of the cCREs (**Fig. 5b, Extended Data Fig. 5c**, **Supplementary Table 4c**). By contrast, DIRs from CTCF binding sites are not enriched for fCREs (**Extended Data Fig. 5d**). We next asked if differentially accessible regions (DARs) are associated with fCREs during differentiation. We analyzed the ATAC-seq data and determined 28,974, 12,246, and 51,101 DARs in pre-differentiation, post-mitotic, and neuronal differentiation, respectively (**Extended Data Fig. 5e**, **Supplementary Table 7**). Notably, fCREs identified from all three screens are enriched for the corresponding DARs (**Fig. 5c, Extended Data Fig. 5f**, **Supplementary Table 4d**), consistent with their role in regulating gene expression in each differentiation stage.

**Figure 5.**
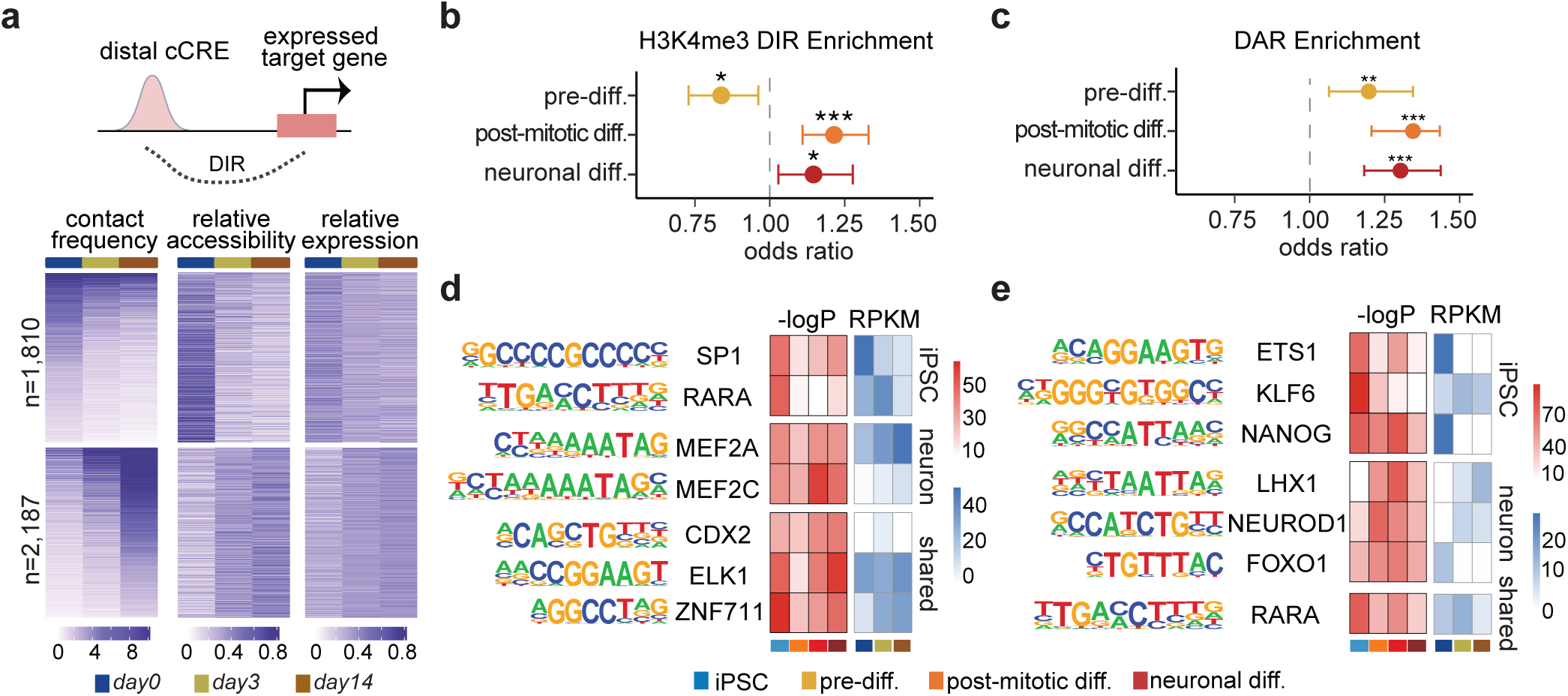
fCREs undergo chromatin remodeling during neuronal differentiation. **a**, Heatmap of XOR neuronal DIR containing a distal accessible distal peak. (Left) The observed H3K4me3 PLAC-seq counts over expected counts. (Middle) Summed normalized ATAC-seq signal within distal 5 Kb bin over total ATAC signal across all timepoints. (Right) Summed TMM-RPKM for all genes within distal 5 Kb bin over total expression across all timepoints. **b-c**, Forest plot showing the association of fCREs with (b) DIRs or (c) DARs. Data are shown as odds ratio with 95% confidence interval (N are indicated in **Supplementary Table 4c-d**, Fisher’s exact test, *: *P* < 0.05, **: *P* < 0.01, ***: *P* < 0.001)**. d-e**, Heatmap showing the top enriched TF motif at TSS (d) or distal (e) fCREs associated with each screen and corresponding TF expression at each time point.

To identify potential TFs that contribute to fCREs function, we determined the TF binding motifs enriched in the promoter-proximal and distal fCREs required for iPSC fitness and neuronal differentiation (**Fig. 5d-e, Supplementary Table 8**). We observed concordant motifs enrichment and TF gene expression in a temporal specific manner along neuronal differentiation. For example, binding motifs for LHX1 and NEUROD1, TFs that are highly expressed in neurons, are enriched at distal fCREs. Interestingly, while FOXO1 binding motif is significantly enriched in fCREs identified in all time points throughout differentiation, the TF itself is only expressed in iPSCs, consistent with its key role in early neuronal fate determination^33^. We also found that binding motifs for key neuronal TFs such as MEF2A, MEF2C are significantly enriched at TSS fCREs while NEUROD1 is significantly enriched at distal fCREs, indicating their distinct roles in gene regulation during neuronal differentiation.

### Neuropsychiatric risk variants are enriched in neuronal fCREs

We next asked whether fCREs are better suited to partition genetic heritability for complex diseases than cCRE annotated based on biochemical signatures alone. We performed linkage disequilibrium score regression^34^ (LDSC) using summary statistics from studies of ten neuropsychiatric disorders^8–11,35–40^. We found that SCZ, ADHD, ASD and post-traumatic stress disorder (PTSD)-associated genetic variants are significantly enriched in neuronal fCREs but not in the non-fitness cCREs (**Fig. 6a**). On the other hand, bipolar disorder (BP) associated variants are enriched in both fCREs and cCREs, while multiple sclerosis (MS) associated variants are exclusively significantly enriched in cCREs, indicating biological processes other than differentiation are associated with these diseases. These results implicate dysregulation of gene expression during neural development as potential contributors of SCZ, ADHD and ASD. By contrast, GWAS variants associated with stroke and Alzheimer’s disease (AD) are not enriched in fCREs or cCREs, suggesting cell types other than neurons are more relevant to these diseases.

**Figure 6.**
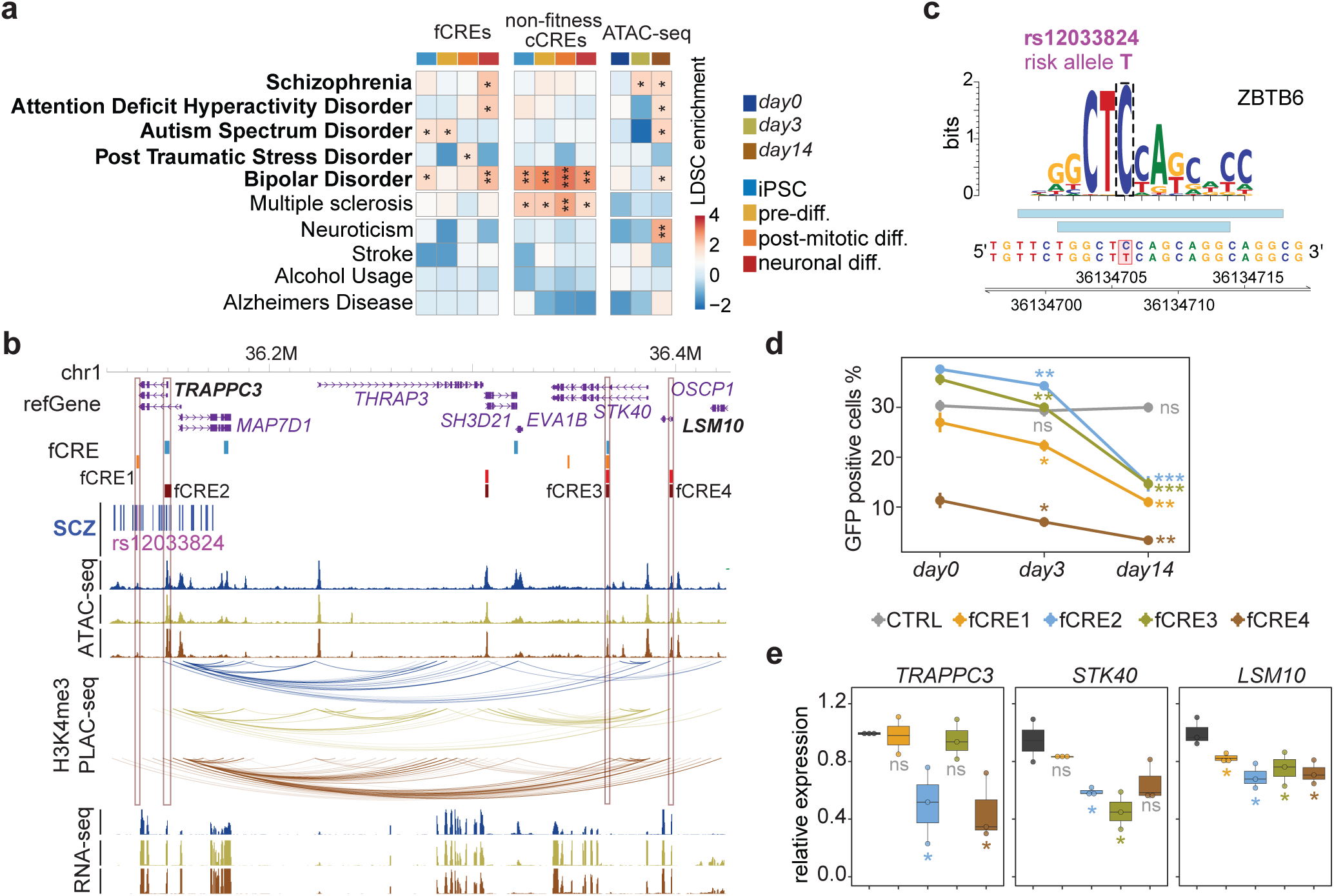
Neuropsychiatric disease-associated risk variants are enriched in neuronal fCREs. **a**, LDSC analysis using annotation of fCREs, non-fitness cCREs or ATAC-seq peaks on neurological disorders. *P*-values are corrected by the Benjamini-Hochberg procedure for multiple tests. FDRs of LDSC coefficients are displayed (*, FDR < 0.05; **, FDR < 0.01; ***, FDR < 0.001). **b**, Genome browser snapshot of the SCZ fine-mapped variants and fCREs near *TRAPPC3*. Purple boxes indicate the validated fCREs. **c**, ZBTB6 motif disrupted by SCZ variant rs12033824 within distal fCRE1 at the *TRAPPC3* locus. **d**, Survival ratio of fCRE perturbed cells by FACS in neuronal differentiation. Data are presented as mean ± s.d. (n = 3, two-tailed two sample t-test). **e**, CRISPRi effect of perturbing fCREs in neurons quantified by RT-qPCRs. Boxplots indicate the median and interquartile range. Whiskers indicate the 5th and 95th percentiles. (n = 3, two-tailed two sample t-test, ns: *P* > 0.05, *: *P* < 0.05, **: *P* < 0.01***: *P* < 0.001).

### Functional analysis of SCZ-associated non-coding variants in fCREs

To explore the roles of neural fCREs in the etiology of SCZ, we first tested the function of fCREs overlapping with several SCZ fine-mapped variants^8^ at the *TRAPPC3*/*LSM10* locus in excitatory neurons (**Fig. 6b**). *TRAPP* is a trafficking protein complex that facilitates vesicle transportation from endoplasmic reticulum to Golgi membrane^41^. Mutations in TRAPP subunits are linked to a wide range of human diseases^42^, including spondyloepiphyseal dysplasia tarda, hypopigmentation, colorectal cancer, and intellectual disability, etc. For example, *TRAPPC3*^43^ and *TRAPPC9*^44^ are SCZ risk genes, and *TRAPPC3* and *LSM10* are known neuronal essential genes at this locus^27^. At this locus, four fCREs are identified by our screens with two of them overlapping with SCZ variants, including rs12033824 and rs12033825 in fCRE1, and rs3754080 and rs10752586 in *TRAPPC3* TSS fCRE2. Notably, the risk allele rs12033824-T could disrupt the binding motif of zinc finger TF ZBTB6, which is highly expressed in neurons and its motif is significantly enriched in neuronal fCREs (**Fig. 6c, Supplementary Table 8**). We found that CRISPRi targeting all four fCREs affect neuronal differentiation (**Fig. 6d, Extended Data Figure 6**). Perturbing fCRE1 resulted in the downregulation of *LSM10*, while suppressing the promoter of either *TRAPPC3* (fCRE2) or *LSM10* (fCRE4) led to simultaneous down-regulation of both *TRAPPC3* and *LSM10* (**Fig. 6e**). These results illustrate the regulatory complexity of disease-associated non-coding variants and suggest both *TRAPPC3* and *LSM10* are regulated by SCZ risk variants in multiple fCREs.

To directly test whether some SCZ non-coding risk variants indeed affect neuronal differentiation, we carried out a high-throughput variant characterization using a newly developed prime editing (PE) screening method^45^. To enable PE screens, we generated a clonal line (i^3^N-nCas9/RT-WTC11) that has a knock-in of an nCas9 and M-MLV reverse transcriptase (RT) expression cassette at the *CLYBL* locus in i^3^N-WTC11 iPSC (**Extended Data Fig. 7a-b**). We chose 110 SCZ-associated variants that were discovered from three large SCZ GWAS studies^8,46,47^ (**Methods**) and were located within 58 fCRE regions, and examined their functions in neuronal differentiation in the PE screen. For each SNP, we designed three pairs of prime editing guide RNA and nick gRNA (pegRNAs/ngRNA) to introduce either reference or alternative allele (a total of 656 pegRNA/ngRNA pairs). We also included 100 non-targeting negative control pegRNA/ngRNA pairs (**Fig. 7a, Supplementary Table 9a**). We infected i^3^N-nCas9/RT-WTC11 iPSCs with a lenti-viral library expressing pegRNA/ngRNA pairs at an MOI of 0.3 with two replicates. Infected cells were selected by hygromycin for 5 days before differentiation to excitatory neurons. To investigate the functional difference between the reference and alternative alleles during neuronal differentiation, we calculated pegRNA copy number fold changes in day-14 neurons versus day-0 iPSCs by MAGeCK^26^. We then obtained the relative fold changes of alternative to reference alleles for each pegRNA/ngRNA at the same nucleotide and identified 45 variants in 29 fCREs as significantly enriched or depleted during neuronal differentiation (FDR < 0.05) (**Fig. 7a**-**b**, **Supplementary Table 9b-c**). 22, 21 and 19 of the 45 functional SNPs were found within the fCREs characterized from the pre-differentiation screen, post-mitotic deafferentation screen and neuronal differentiation screen, suggesting different roles of functional variants in controlling neural developmental genes in SCZ (**Fig. 7c**). For example, rs1801311 is located in the first exon of *NDUFA6*, which has been implicated in SCZ risk with regulatory functions^8^, The reference allele rs1801311-G, which is also a SCZ risk allele, could disrupt the binding motifs of POLAR2A, TAF1, and YY1, leads to reduced enhancer activity in luciferase reporter assays^48^, and upregulates *NAGA* expression in the SCZ patient brain^49^. Our study further provides direct evidence that the rs1801311-G allele can confer SCZ risk by reducing neuronal differentiation capacity by 75% (Alt/Ref log_2_(fold change) = 2.03) compared to the A allele.

**Figure 7.**
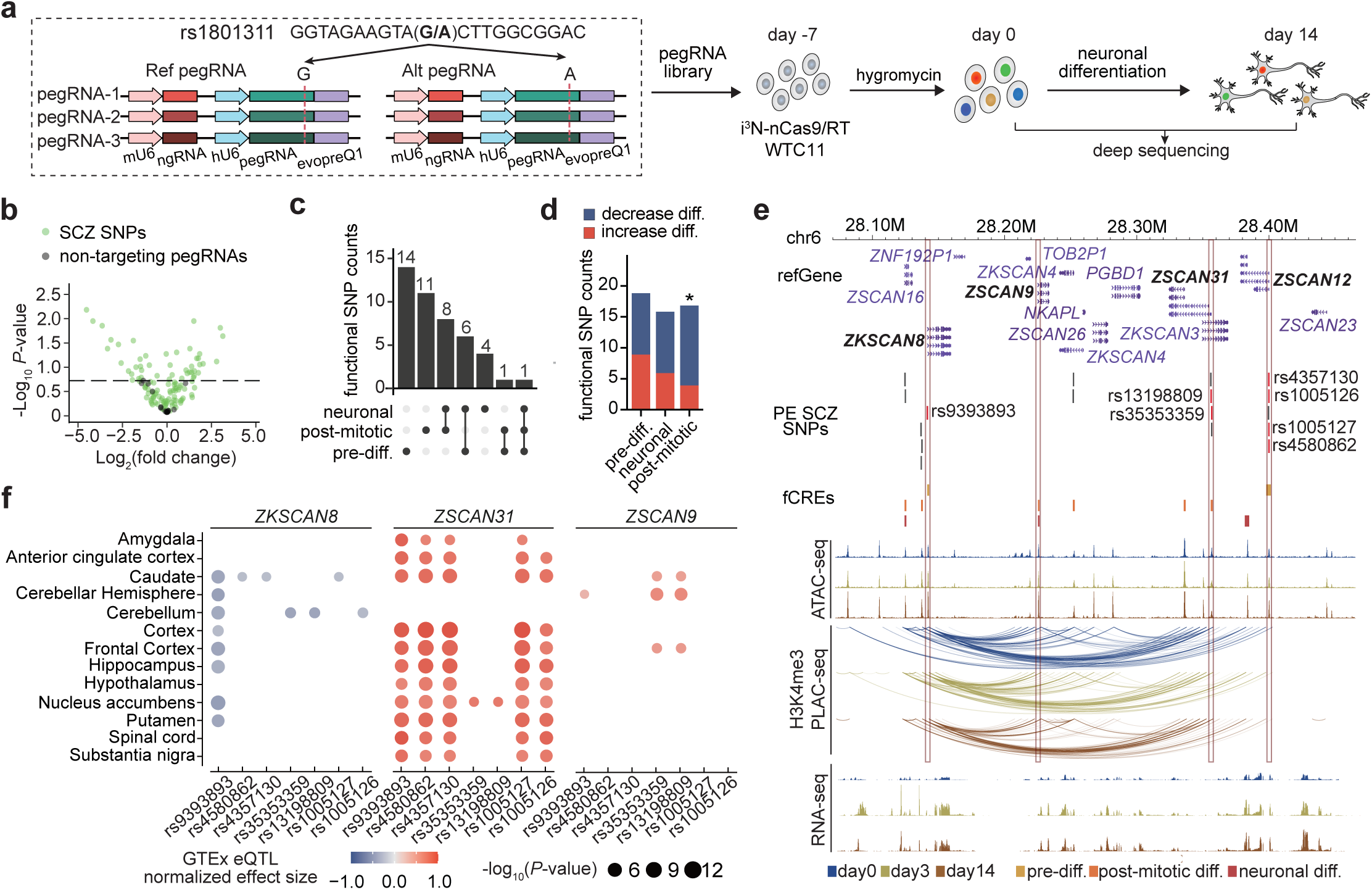
Functional characterization of SCZ-risk variants within the fCREs. **a**, (Top) Design of pegRNAs for SCZ-related SNPs. For each variant, three replicates of pegRNA/ngRNA pairs were designed to introduce either the Alt or Ref allele. (Bottom) Workflow of the PE screen. **b**, Volcano plot illustrating the SNPs’ impact on neuronal differentiation, with positive or negative effects determined by the relative effect of the Alt versus Ref allele. The dashed line represents the *P* value cutoff determined by non-targeting pegRNAs (FDR < 0.05). n = 2. **c**, Upset plot demonstrating the shared functional SNPs across different types of fCREs. **d**, Bar plot illustrating the effects of risk alleles of functional SNPs on different stages of neuronal differentiation. *P*-values were calculated using the two-sided binomial test. (*: *P* < 0.05) **e**, Genome browser snapshot showing a region with the SCZ risk variants and fCREs enriched with multiple SCAN domain-containing TFs. Red-labeled SNPs indicate positive functional SNPs. **f**, Normalized effect size and *P* values for functional SNPs on regulating target genes based on the GTEx eQTL data in different brain regions. Colors represent normalized effect size and sizes represent *P* values.

Interestingly, we found divergent effects of SCZ risk variants at different stages of neuronal differentiation based on specific differentiation stages. Specifically, risk alleles are more likely to negatively impact post-mitotic differentiation stage (two-sided exact binomial test, *P* = 0.049) than other stages (**Fig. 7d**), consistent with disrupted neurogenesis in SCZ patient derived organoids compared to normal organoids^50^.

34 of the 45 functional variants are located in the promoters of 21 genes, and 4 of them, rs2524093, rs2524092, rs2844622 and rs28626310, are located in the promoter of *HLA-C* gene (**Extended Data Fig. 7c**). While HLA-C has been recognized for its involvement in immune cell types, recent studies have highlighted the potential involvement of neuronal MHC-I, including HLA-C, in neurodevelopment and neurodegenerative diseases^51,52^. *HLA-C* expression is transiently increased at day 3 after differentiation compared to day 0 and day 14. GTEx^53,54^ eQTL data demonstrated a correlation between three SNPs (rs2524093, rs2524092 and rs2844622) and *HLA-C* expression in brain tissues, with risk alleles associated with increased *HLA-C* expression. It is worth noting that the *HLA-C* promoter interacts with other gene promoters. However, none of these interacting genes are known essential genes or TSS fCREs, suggesting that these four SNPs may exert their functional effects by directly regulating *HLA-C*. Our analyses thus expand the understanding of *HLA-C* beyond its classical immune-related functions and emphasize its potential contributions to neurodevelopmental disorders.

Seven functional variants are identified in the locus enriched with multiple SCAN domain-containing TFs, which include rs9393893 in the promoter of *ZKSCAN8*, rs13198809, rs35353359 in the promoter of the *ZSCAN31*, and rs1005126, rs4580862, rs4357130, and rs1005127 in the promoter of *ZSCAN12*. All three genes are neuronal genes, with *ZKSCAN8* and *ZSCAN12* in pre-differentiation, and *ZSCAN31* in post-mitotic differentiation (**Fig. 7e**). These three promoters interact with each other, and they also interact with the promoter of another essential gene, *ZSCAN9,* in post-mitotic differentiation and neuronal differentiation. The expression of three ZSCAN genes, *ZKSCAN8, ZSCAN12 and ZSCAN9* are upregulated in day 3 and day 14 differentiated cells (**Fig. 7e**), consistent with their role in neuronal differentiation. GTEx^53,54^ eQTL data demonstrates that these seven SNPs are associated with the expression of *ZKSCAN8*, *ZSCAN31*, and *ZSCAN9* from different brain regions, highlighting the complexity of this locus with disease-associated variants cooperatively affecting multiple genes through both promoter and enhancer regulations (**Fig. 7f**). In summary, our prime editing screening provides a deeper understanding of the functional SNPs located in fCREs at the base pair resolution. Importantly, our approach offers more detailed functional insights into the role of SCZ SNPs during neuronal differentiation.

## Discussion

In this study, we performed genome-scale CRISPRi screens to interrogate the function of 22,000 cCREs in iPSC undergoing neuronal differentiation. We identified 2,857 fCREs required for iPSC fitness and 5,540 fCREs required for excitatory neuron differentiation. We delineated differential chromatin features between fCREs and non-essential cCREs, observing increased chromatin accessibility, stronger active H3K27ac and H3K4me3 signals, and a greater quantity of chromatin interactions at fCREs. Furthermore, neuronal fCREs exhibited both increasing chromatin accessibility and interactions over the neuronal differentiation time course. To our knowledge, this study represents the most extensive functional characterization of cCREs in human neurons to date, expanding our knowledge of essential genes^17,18,20,27^ and functional regulatory sequences regulating iPSC fitness and neuron differentiation. The annotation of fCREs provides a great resource of essential DNA elements and genes, and the identified TF binding motifs within promoter-proximal and/or distal fCREs bring new biological insights into iPSC fitness and neuronal differentiation.

Individual CRISPRi validations targeting *MYC* and *SOX2* fCREs in iPSCs and *TUBA1A* and *EPHB1* fCREs in neurons not only confirm the fCREs regulatory role in essential gene expression, but also exhibit diverse gene regulatory models including promoter-promoter interactions^55,56^ and co-regulation of multiple target genes^57^. Our assays also confirmed the functionalities of 113 out of 282 developmental brain enhancers previously characterized in transgenic mouse assays^22^.

Previous studies have demonstrated that neuropsychiatric genetic variants are enriched in neuronal cCREs annotated by biochemical signatures^12–14^. In our study, we further narrow-down cCREs to functionally validated fCREs, which offers a better pathway for interpreting the non-coding risk variants associated with neuropsychiatric diseases. In particular, we revealed that fCREs accounts for a high degree of genetic heritability for neurodevelopmental diseases including SCZ, ASD, and ADHD, compared to cCREs, underscoring the value of identifying fCREs during the neuronal differentiation. We further explored SCZ fine-mapped variants using multiple approaches. We first investigated two fCREs covering SCZ variants at the *TRAPPC3*/*LSM10* locus, and found that perturbations on SCZ risk fCREs affects neuronal differentiation and *TRAPPC3*, *LSM10* gene expressions. We further performed a prime editing screen targeting 110 SCZ genetic variants within neuronal fCREs during neuronal differentiation. 45 causal variants in 29 fCREs were identified as essential for the neuronal differentiation process. These functional validations are consistent with GTEx eQTL datasets at the molecular level. Our findings also suggest that SCZ risk genes including *HLA-C*, *ZKSCAN8*, *ZSCAN9*, *ZSCAN12* and *ZSCAN31* may contribute to disease etiology by affecting neuronal development. These results confirmed the significance of functional characterization of neuronal fCREs in understanding the role of neuropsychiatric genetic variants, and identified causal disease risk variants. As the first high-throughput screen at base-pair level for SCZ variants in neurons, our prime-editing screen results provided a paradigm of characterizing endogenous function of GWAS variants in relevant cell types at scale.

fCREs in our study are identified based on fitness and differentiation phenotypes. Non-essential cCREs in current screens could be functional in other cell types and biologically processes. Future studies employing other cell types or neuronal phenotypes such as calcium dynamics and synaptic connectivity should offer additional annotation for fCREs and functional disease-associated variants.

## Acknowledgements

This work was supported by the National Institutes of Health (NIH) grants UM1HG009402 (to Y.S. and B.R.), and U01DA052713 (to Y.S.). M.H. was partially supported by the NIH grants R35HG011922 and UM1HG011585. J.W. is supported by T32ES007018. This work was made possible in part by NIH grants P30DK063720, and S101S10OD021822-01 to the UCSF Parnassus Flow Cytometry Core. Sequencing was performed at the UCSF CAT, supported by UCSF PBBR, RRP IMIA, and NIH 1S10OD028511-01 grants.

## Author contributions

Y.S. and B.R. conceived the study. Y.S. and B.R. supervised the study. X.Y., I.R.J., P.B.C., H.Y., X.R., C.B. and X.C. performed experiments under the supervision of Y.S. and B.R.. X.Y., I.R.J., P.B.C., H.Y., L.Z., B.L., Y.E.L., Q.S. and J.W. performed computational analysis under the supervision of Y.S., M.H., Y.L. and B.R.. X.Y., I.R.J., P.B.C., H.Y., M.H., B.R. and Y.S. analyzed and interpreted the data. X.Y., I.R.J, P.B.C., H.Y., B.R., and Y.S. prepared the manuscript with input from all other authors.

## Competing interests statement

B.R. is a co-founder and consultant of Arima Genomics Inc. and co-founder of Epigenome Technologies. X.R., H.Y., and Y.S. have filed a patent application related to pooled prime editing screens. The other authors declare no competing interests.

## Code availability statement

Data are analyzed using published pipelines with parameters described in the Methods section. No custom code is developed in this study.

## Data availability statement

PLAC-seq, RNA-seq, and ATAC-seq datasets used in this study are available at the Gene Expression Omnibus under the accession number GSE236705 (reviewer token: cvsbsekirhartud). CRISPRi screens in iPSC and neurons are released by ENCODE portal under the functional characterization experiment series ENCSR499VAT and ENCSR737MRW. Data can be visualized on the WashU Epigenome Browser using the following session bundle ID: 00a68230-389c-11ee-840b-9392e91a6ccc.

## Supplementary Tables

Supplementary Table 1 cCRE prioritization and gRNA design in iPSC and neuron CRISPRi screen

Supplementary Table 2 MAGeCK analyses for fCREs in iPSC and neuron CRISPRi screens

Supplementary Table 3 Epigenetic profiling data processing metrics in neuronal differentiation.

Supplementary Table 4 Association analysis for fCREs and chromatin loops.

Supplementary Table 5 GO enrichment analysis for essential genes and distal fCRE target genes annotated by H3K4me3-or CTCF-associated interactions

Supplementary Table 6 DIRs along neuronal differentiation

Supplementary Table 7 DARs along neuronal differentiation

Supplementary Table 8 Motif analysis for TSSs and distal fCREs in iPSCs and neurons

Supplementary table 9 PegRNA design, counts and functional SCZ SNPs in the prime editing screen

Supplementary Table 10 RT-qPCRs primers in individual CRISPRi experiments

## Methods

### Ethics statement

The use of the iPSC line WTC11^58^ was approved by the Human Gamete, Embryo and Stem Cell Research (GESCR) Committee at UCSF.

### Cell culture

iPSCs were cultured with mTesR on Matrigel coated plates. Excitatory neurons were derived from isogenic and inducible neurogenin-2 (Ngn2) WTC11 iPSCs (i^3^N-WTC11) as previously described^15^. i^3^N-WTC11 iPSCs stably expressing dCas9-KRAB or nCas9/RT were generated according to an established method^59^. dCas9-KRAB or nCas9/RT driven by CAG promoter was knocked-in to a safe harbor locus in the intronic region of *CLYBL* to enable robust transgene expression throughout differentiation. Excitatory neurons were derived in a simplified two-step protocol. In the pre-differentiation stage, i^3^N-WTC11 iPSCs were cultured with medium containing knockout DMEM/F12 supplemented with 1× N-2, 1× NEAA, 1 μg/ml mouse laminin, 10 ng/ml brain-derived neurotrophic factor (BDNF), 10 ng/ml NT3 and 2 μg/ml doxycycline. ROCK inhibitor (10 μM) was added only for the first day. Medium was changed daily for 3 days. In the post-mitotic differentiation stage, neural progenitor cells were dissociated by Accutase and re-seeded onto Poly-L-ornithine coated plates in medium containing equal parts DMEM/F12 and Neurobasal-A supplemented with 0.5× B-27, 0.5× N-2, 1× NEAA, 0.5× GlutaMax, 1 μg/ml mouse laminin, 10 ng/ml BDNF, 10 ng/ml NT3 and 2 μg/ml doxycycline. The doxycycline was omitted in all subsequent medium changes. Half of the medium was changed on day 7 and 2-week excitatory neurons were used for CRISPRi screens.

### Design and assembly of the CRISPRi gRNA libraries

iPSC and neuronal essential genes were selected based on previous publications^17,18,20,27^. 18,220 cCREs in iPSCs and 14,736 cCREs in neurons were prioritized as described in the main text for gRNA design. Neural VISTA enhancers were selected if the enhancers are active in neural tube, forebrain, midbrain, hindbrain, dorsal root ganglion and cranial nerve. gRNAs were designed by CRISPR-SE^60^ and selected based on the cutting specificity score greater than 0.3 from GuideScan^61^ (v2.0.6). Top 10 ranked gRNA were selected and dual gRNAs were paired by maximizing their distance within each cCREs. 17,743 in iPSC and 14,411 cCREs in neurons were successfully targeted in gRNA design. For negative controls we randomly selected negative gRNAs from previous genome-scale screens^24,25^. 8,389 safe targeting gRNAs and 1,011 non-targeting gRNAs were included in the screen. The final library contains 88,715 and 66,652 pairs of gRNAs for iPSC and neuron individually. Dual gRNA oligos were synthesized by Agilent and plasmid libraries were assembled in two steps as previously described^56^. In step 1 we amplified the oligo libraries by PCR and purified them with AMPure XP beads. We then performed Gibson assembly reactions to insert gRNA pools into the lenti-Guide-puro backbone (Addgene 52963). The purified product was electroporated into Endura competent cells (Lucigen). We extracted the plasmid pool using the Plasmid Maxi prep kit. In step 2 we digested plasmid pools in step-1 at BsmB-I cutting site between the synthesized paired gRNAs, and ligated with the DNA fragment containing gRNA-scaffold, a linker fragment and the mouse-U6 promoter. The purified DNA ligation product was electroporated into Endura competent cells and plasmid pools were extracted as the final library. Coverage of the library was tested by paired gRNA matches based on amplicons covering the human-U6 promoter and the second gRNA-scaffold. Each step we used a coverage of 1,000 transduced cells per dual-gRNA pair. After library assembly cCREs targeted by low-abundance gRNA pairs were excluded. In total we successfully tested 16,670 cCREs in iPSC and 14,289 cCREs in neuronal CRISPRi screenings.

### Pooled CRISPRi screen for iPSC proliferation and neuronal differentiation

Library plasmid pools were packaged into lenti-virus by PsPAX2 (Addgene 12260) and pMD2.G (Addgene 12259) in 293T cells. Lentivirus were concentrated by high-speed centrifuge and titered in iPSC using the survival ratio under antibiotics selection from a serial dilution of lentivirus transduction. In the screens, iPSCs were infected at a low multiplicity of infection (MOI = 0.5) to ensure each infected cell got only one viral particle. 24hrs after infection cells were re-seeded with low density and underwent puromycin selection (0.5 mg/mL, Gibco A1113803) for pooled CRISPRi screen for 5 days. On day 7 after lenti-viral infection we collected cells with a coverage of 1,000 transduced cells per dual-gRNA pair as day-0 library. For the iPSC screen, we kept culturing the cells in regular medium for another 14 days and then collected cells as day-14 library. For the neuronal screen, we started pre-differentiation after antibiotic selection, and collected cells 3 days later after pre-differentiation and another 11 days later after post-mitotic differentiation also with a coverage of 1,000 transduced cells per dual-gRNA pair.

### Genome wide CRISPRi screen data analysis

The abundance of dual-gRNA pairs from different time points was mapped to the initially designed oligo library sequences using BWA^62^ (v0.7.17). We used MAGeCK^26^ (v0.5.9.4) to identify fCREs required for cell proliferation. Non-targeting dual-gRNA pairs were provided to generate the null distribution when calculating the *P* values. dual-gRNA pairs were ranked based on the *p*-values, and we used the modified RRA algorithm from MAGeCK to obtain the RRA score for the individual target gene or cCRE. The threshold value for candidate fCREs is the empirical FDR < 0.05 based on the percentage of candidate dual-gRNA pairs identified from negative controls above this threshold, including non-essential genes and safe-target control genomic regions to account for the possible toxicities from the silencing mediated by dCas9-KRAB.

### Annotating fCREs

TSS fCREs were defined as fCREs overlapping with a transcriptionally active TSS with an RPKM > 1 at any time point of day 0, day 3, or day 14. Distal fCREs were defined as fCREs that do not overlap with any TSSs or overlap with TSSs with RPKM < 1 at all timepoints.

### CRIPSRi validation followed by RT-qPCR or FACS

gRNA targeting individual fCREs were synthesized and inserted into modified lenti-Guide puro with GFP tag. iPSCs were infected with lentivirus at MOI ∼1. To detect cell survival using FACS, cells were not selected by puromycin but cultured in regular medium for 3 days to ensure all infected cells showed positive GFP expression. We then did FACS analysis to count GFP positive cells on day 0 and day 14, plus day 3 for neurons. Contour plots for gating and representative dot plots are summarized in **Extended Data Fig. 6**. To detect target gene expression by RT-qPCR, cells were selected with puromycin for 5 days. Then we kept culturing iPSC for 14 days before mRNA was extracted. Meanwhile we started neuronal differentiation and mRNA was extracted after 14 days. RT-qPCR primers are listed in **Supplementary Table 10**.

### RNA-seq

RNeasy Plus Mini Kit (Qiagen 74134) was used to isolate total RNA and approximately 1 ug of RNA was processed with the TruSeq Stranded mRNA Library Prep Kit (Illumina 20020594) to prepare libraries for paired-end sequencing on Nova-seq 6000. Raw paired-end fastq RNA-seq libraries were filtered for high quality reads and trimmed to 100bp using fastp (v0.22). Next, the libraries were aligned to hg38 using STAR (v2.7.10a) running the standard ENCODE parameters and strand-specific quantification was performed using RSEM (v1.2.28) with the GENCODE 38 annotation. TMM-normalized RPKM values for each gene were obtained via the edgeR package. The mean values across all replicates were used for all downstream analyses.

### ATAC-seq

ATAC-seq was performed using the Nextera DNA Library Prep Kit (Illumina FC-121-1030). Briefly, each sample received a 1x wash with ice-cold 1x PBS followed by resuspension in ice-cold nuclei extraction buffer (10 mM Tris-HCl pH 7.5, 10 mM NaCl, 3 mM MgCl_2_, 0.1% Igepal CA630, and 1x protease inhibitor) for 5 min. 50,000-100,000 cells per reaction were subsequently transferred into 50 μL 1x buffer TD with 2.5 μL TDE1 enzyme for 30 min at 37 °C for transposition. DNA was purified with Qiagen MinElute spin columns (Qiagen 28006), PCR amplified, and size-selected with AMPure XP beads. Libraries were then paired-end sequenced on Nova-seq 6000. Reads were trimmed to 50 bp and filtering for high quality reads with fastp (v0.22) were then mapped to hg38 and processed with the ENCODE pipeline (v1.10.0) with default settings. Optimal IDR peaks for each timepoint were used for all downstream analysis. To call DARs, bam and narrowpeak files for each timepoint were supplied to the R package Diffbind^63^ (v3.4.11) and the DEseq2^64^ (v1.38.3) method was used to identify differential peaks, FDR < 0.05 and log_2_(fold change) < 0.5, from a union peak set.

### PLAC-seq

PLAC-seq libraries were generated using the Arima-HiC^+^ kit (P/N A101020) according to the manufacturer’s protocol. Briefly, 3-5 million cells or approximately 15 ug of DNA crosslinked with 2% PFA (Fisher Scientific F79-500) were used as input per reaction. Following restriction digestion, biotinylation, and proximal ligation, the chromatin was sheared with a Covaris S220 sonicator to fragments between 300 and 1,000 bp and subsequently and immunoprecipitated with anti-H3K4me3 antibody (Millipore 04-745) or anti-CTCF antibody (Active Motif 91285). Immunoprecipitated chromatin was then indexed with the Accel-NGS 2S Plus DNA Library Kit (Swift Biosciences 21024) and amplified with KAPA Library Amplification Kit (Roche KK2620), before paired-end next generation sequencing.

### PLAC-seq data processing

We used MAPS^65^ (v1.1.0) to call significant interactions using default parameters at 5 Kb resolution for both H3K4me3 and CTCF associated PLAC-seq libraries. Briefly, BWA (v0.7.17) was used to map the raw reads to hg38. After removing low quality and non-mapped reads, anchor regions were defined by using the MACS2 (v2.2.4) callpeak command with the following options “--broad --nolambda --broad-cutoff 0.01” on read pairs with an interaction distance < 1 Kb normalized by sequencing depth for each timepoint. A final anchor set was generated by taking the union of all timepoints for both H3K4me3 and CTCF conditions. Next read pairs were defined as AND, XOR, or NOT interactions based on whether both, one, or neither of the contact bins overlapped with union anchors. Only intrachromosomal AND and XOR pairs between 10 Kb ∼ 2 Mb were retained for downstream analysis. HPrep^66^ (v1.0) was used to confirm reproducibility between replicates before merging them. Significant interactions were called using approximately 41 ∼ 43 million usable reads per time point. Significant interactions were called as previously described^65^.

### Contact Probability

We calculated contact probability similarly as previously described^31^. Briefly, we first determined the total number of paired PLAC-seq reads between 10 Kb and 2,560 Kb. Next, we obtained the contact probability as the sum of observed paired counts within a log2 distance divided by the total. Finally, we plotted the best fitting line between distances using a loess curve.

### Epigenomic Signal Comparison

H3K27ac (ENCFF573DHK), H3K4me3 (ENCFF577IZN) and CTCF (ENCFF974CGC) ChIP-seq signals were obtained from ENCODE, while the ATAC-seq and PLAC-seq contacts were generated in this study. Next, the bigWigAverageOverBed (v302.1) command was used to obtain the signal overlapping with the 16,583 cCREs in the IPSC screen for the H3K27ac, H3K4me3, CTCF and ATAC signal. The bedtools (v2.29.0) pairToBed command was used to count the number of interactions per cCRE. Finally, the Wilcoxon test measured significant differences between the mean values.

### Target genes annotation for distal fCREs and GO enrichment analysis

Target genes of distal fCREs were annotated by PLAC-seq datasets. First, anchor bins are annotated by TSS overlap. Then distal fCREs were intersected with both anchor bins and target bins to get all potential target genes (**Supplementary Table 5**). We further filtered target genes by RPKM > 1 in day 0 for iPSCs, in day 3 or day 14 for neurons, and excluded genes that have been validated as non-essential in our screens. GO enrichment analysis was performed using clusterProfiler^67^ (v3.18.1). Known essential genes, new essential genes and distal fCRE target genes annotated by H3K4me3-and CTCF-associated PLAC-seq were used as input compared to a genome wide background gene list. Enriched GO terms in the “GO Biological Process” ontology with q-value < 0.05 were reported (**Supplementary Table 5**).

### Identification of DIRs

We applied edgeR^68^ (v3.32.1) to identify DIRs as described in our previous study^69^. We used the comparison between H3K4me3 PLAC-seq data at day 0 and day 3 as an illustrative example to demonstrate our data analysis procedure. Briefly, we first used MACS2^70^ (v2.2.4) to call H3K4me3 ChIP-seq peaks, described above, at day 0 and day 3, and defined differential H3K4me3 ChIP-seq peaks between two time point if (1) FDR < 5% and (2) log_2_(fold change) > 0.5. Next, we selected 5 Kb bin pairs anchored at the non-differential H3K4me3 ChIP-seq peaks for the differential chromatin interaction analysis, to ensure that the dynamics of chromatin contact frequency is not biased by the dynamics of protein binding intensity. We started from 5 Kb bin pairs with 1D genomic distance 5 Kb ∼ 2 Mb, and contained >= 20 reads in at least one replicate. We then stratified all selected 5 Kb bin pairs into 1D genomic distance groups from 5 Kb to 150 Kb. For the remaining 5 Kb bin pairs with 1D genomic distance > 150 Kb, we stratified them into groups such that each group contains the same number of 5 Kb bin pairs as that in the group of 5 Kb bin pairs with 1D genomic distance 150 Kb. For 5 Kb bin pairs in each group, we applied edgeR to evaluate the statistical significance of the difference in chromatin contact frequency. We defined a 5 Kb bin pair as a DIR if FDR < 0.01 and log_2_(fold change) > 1. To visualize the identified DIRs, we plotted 1D genomic distance vs. the magnitude of change of chromatin contact frequency, defined as the signed -log_10_(*P*-value). We applied the same approach to identify DIR for H3K4me3 PLAC-seq data between day 0 and day 14, day 3 and day 14, and CTCF PLAC-seq data between day 0 and day 14.

### Heatmap

Significant interactions specific to one time point or interactions labeled as DIRs between two timepoints were prioritized. Next, we further filtered for only XOR interactions. The contact frequency was determined by the observed count between bin pairs over the expected count generated by MAPs. Next the ratio of ATAC-seq signal was calculated in the following manner. First, a union peak set of all time points was made by concatenating all IDR peaks and subsequently merging them with the bedtools merge command. Next, the ATAC-seq signal for each time point was obtained by using the bigWigAverageOverBed from sequencing depth normalized bigwig files. The signal was further quantile-normalized with the normalize.quantiles() function in R (v4.2.2). Lastly, if multiple signals occurred within the same bin, they were summed. The heatmap displays the percentage of the individual time point over the total signal summed across all samples. Similarly, transcription of all genes within the anchor bin was displayed as the individual TMM-RPKM over the total expression summed across all samples.

### Motif analysis

Motif enrichment analysis using HOMER^71^ (v4.11.1) running the default settings. fCREs from iPSC screen and pooled neural screens were used as input. ATAC-seq peaks in iPSC and day 14 neurons were used as background. The hypergeometric distribution was used for motif scoring. Significance and expression values for top detected motif and its corresponding TFs are reported. Cell type specific or shared motifs are categories based on motif enrichment lists, with the cell type specific TF expression considered. A full list of motifs is reported in **Supplementary Table 8**.

### LDSC analysis

Three sets of genomic regions were used for LDSC (v1.0.1) analyses, including 1) fCREs from CRISPRi screens; 2) non-fitness cCREs from CRISPRi screens. 3) the chromatin accessible regions were identified from ATAC-seq for day 0 iPSC, day 3 and day 14 neurons. To enable comparison to GWAS of human phenotypes, we used liftOver with default setting “-minMatch=0.95” to convert accessible elements from hg38 to hg19 genomic coordinates. GWAS summary statistics were downloaded for quantitative traits related to neurological disease including SCZ^8^, ADHD^9^, ASD^10^, BD^11^, PTSD^35^, MS^36^, neuroticism^37^, stroke^38^, alcohol usage^40^ and AD^39^. We prepared summary statistics to the standard format for LD score regression^34^. For the analysis using fCREs, non-fitness cCREs, and ATAC-seq peaks as annotations, we take the superset of chromatin accessible regions in day 0 iPSC, day 3 and day 14 neurons as the background control. For each trait, LD score regression was performed to estimate the enrichment coefficient of each annotation jointly with the background controls.

### Design and assembly of the prime editing screening library

SCZ GWAS SNPs were initially selected if supported by at least two studies from four SCZ studies^8,46,47,72^. A significance threshold of *P* < 5e-8 was applied to the three GWAS studies^8,46,47^, while for DeepGWAS^72^, a posterior probability greater than 0.5 was utilized. SNPs in linkage disequilibrium with these initial variants (r^2^ > 0.6) were also incorporated from the TOP-LD database^73^, focusing on the EUR population, resulting in a comprehensive collection of 25,999 SNPs. Subsequently, we identified 110 variants situated within neuronal differentiation fCREs. Prime editing was performed using PE3 system^74^. For each SNP, pegRNAs were designed to edit the variants into either reference or alternative alleles using the default parameters of the PrimeDesign software^75^ (v0.2). To increase transcription efficiency from the U6 promoter, a guanine nucleotide was added to the 5’ end of all pegRNAs/ngRNAs with leading nucleotides other than G. We eliminated pegRNA/ngRNA pairs containing BsmBI sites (GAGACG, CGTCTC) or a TTTTT sequence in the pegRNA spacer, ngRNA spacer, or pegRNA extension. When multiple pairs were available for the same locus, pegRNA/ngRNA pairs were selected to maximize specificity, efficiency, and the distance between ngRNA and pegRNA, while minimizing the editing distance between pegRNA and the target. For each allele, we attempted to design three unique pairs of pegRNAs. For non-targeting pegRNA/ngRNA pairs, the pegRNA spacer, ngRNA spacer, and pegRNA extension sequences were selected from the ENCODE non-targeting sgRNA reference dataset (https://www.encodeproject.org/files/ENCFF058BPG/). The final pegRNA pool comprised 656 experimental pegRNAs and 100 negative control pegRNAs that do not target the human genome. To link these component sequences, we used the following template: 5’-CTTGGAGAAAAGCCTTGTTT[ngRNA-spacer]GTTTAGAGACG[5nt-random-sequence]CGTCTCACACC[pegRNA-spacer]GTTTTAGAGCTAGAAATAGCAAGTTAAAATAAGGCTAGTCCGTTATCAACTTGAAAAAGTGGCACCGAGTCGGTGC[pegRNA-extension]CCTAACACCGCGGTTC-3’.

Library oligos for the prime editing screen were synthesized by Twist Bioscience and amplified using the NEBNext High-Fidelity 2× PCR Master Mix (NEB M0541L) with the forward primer GTGTTTTGAGACTATAAATATCCCTTGGAGAAAAGCCTTGTTT and the reverse primer CTAGTTGGTTTAACGCGTAACTAGATAGAACCGCGGTGTTAGG. To amplify paired PegRNA/ngRNA library oligos, we employed emulsion PCR (ePCR) to reduce recombination of similar amplicons during PCR. In brief, we performed 96 20 μl ePCR reactions using 0.01 fmol of pooled oligos with NEBNext High-Fidelity 2× PCR Master Mix. Each 20 μl PCR mix was combined with 40 μl of an oil-surfactant mixture (containing 4.5% Span 80 (v/v), 0.4% Tween 80 (v/v), and 0.05% Triton X-100 (v/v) in mineral oil). This mixture was vortexed at maximum speed for 5 min, briefly centrifuged, and placed into the PCR machine for amplification. The thermocycler settings were as follows: 98 °C for 30 s, followed by 26 cycles (98 °C for 10 s, 60 °C for 20 s, 72 °C for 30 s), then 72 °C for 5 min, and finally a 4 °C hold. The ramp rate for each step was 2°C/s. After PCR, individual reactions were combined and purified using the QIAQuick PCR Purification Kit (Qiagen 28104). Purified PCR products were then treated with Exonuclease I (NEB M0568L) and purified using 1× AMPure XP beads (Beckman Coulter A63881). The isolated ePCR products were inserted into a BsmBI-digested lentiV2-mU6-evopreQ1 vector via Gibson assembly (NEB E2621L). The assembled products were electroporated into Endura electrocompetent cells (Biosearch Technologies 60242), and approximately 4,000 independent bacterial colonies were cultured for each library. The resulting plasmid DNA was linearized by BsmBI digestion, gel-purified, and ligated using T4 ligase (NEB M0202M) to a DNA fragment containing an sgRNA scaffold and the human U6 promoter. The resulting library was electroporated into Endura electrocompetent cells and cultured as described above. Finally, the plasmid library was extracted using the Qiagen EndoFree Plasmid Mega Kit (Qiagen 12381).

### Prime editing screen data analysis

Sequencing libraries underwent an initial step of trimming 5 bp random sequences from both read-1 and read-2. Subsequently, low-quality reads were filtered out using fastp (v0.22) before formal mapping. Read counts were calculated based on specific criteria for each pegRNA/ngRNA pair, which included: (1) read-1 exactly matches the sequence containing a 20-21 nt ngRNA spacer and 5 bp flanking sequences, and (2) read-2 exactly matches the reverse complementary sequence containing the complete pegRNA extension and 5 bp flanking sequences. For all samples, oligo counts were inputted into MAGeCK^26^ (v0.5.9.4) to determine the fold changes of each oligo by comparing the day 14 data to the day 0 data with the normalization with control non-targeting. Subsequently, the log2(fold change) of each pegRNA was used to calculate the relative fold change for SNPs (alternative to reference). In order to create the control non-targeting SNP, we randomly grouped two pegRNA/ngRNA designs as reference and alternative for the non-targeting SNPs, and three pairs were grouped together for a single non-targeting SNPs design. To calculate the fold change and *P*-value of the SNPs, we combined all experimental SNPs and non-targeting SNPs and utilized the MAGeCK (v0.5.9.4) RRA method. To minimize false positive results and maintain an empirical FDR below 5%, we selected a *P*-value cutoff corresponding to the minimal *P*-values obtained from the non-targeting SNPs.

### Statistics and reproducibility

Statistical methods used are indicated in the figure legends or Methods sections. No statistical method was used to predetermine sample size, and no data were excluded from analysis. The experiments were not randomized. The Investigators were not blinded to allocation during experiments and outcome assessment.

---

**Extended Data Figure 1.**
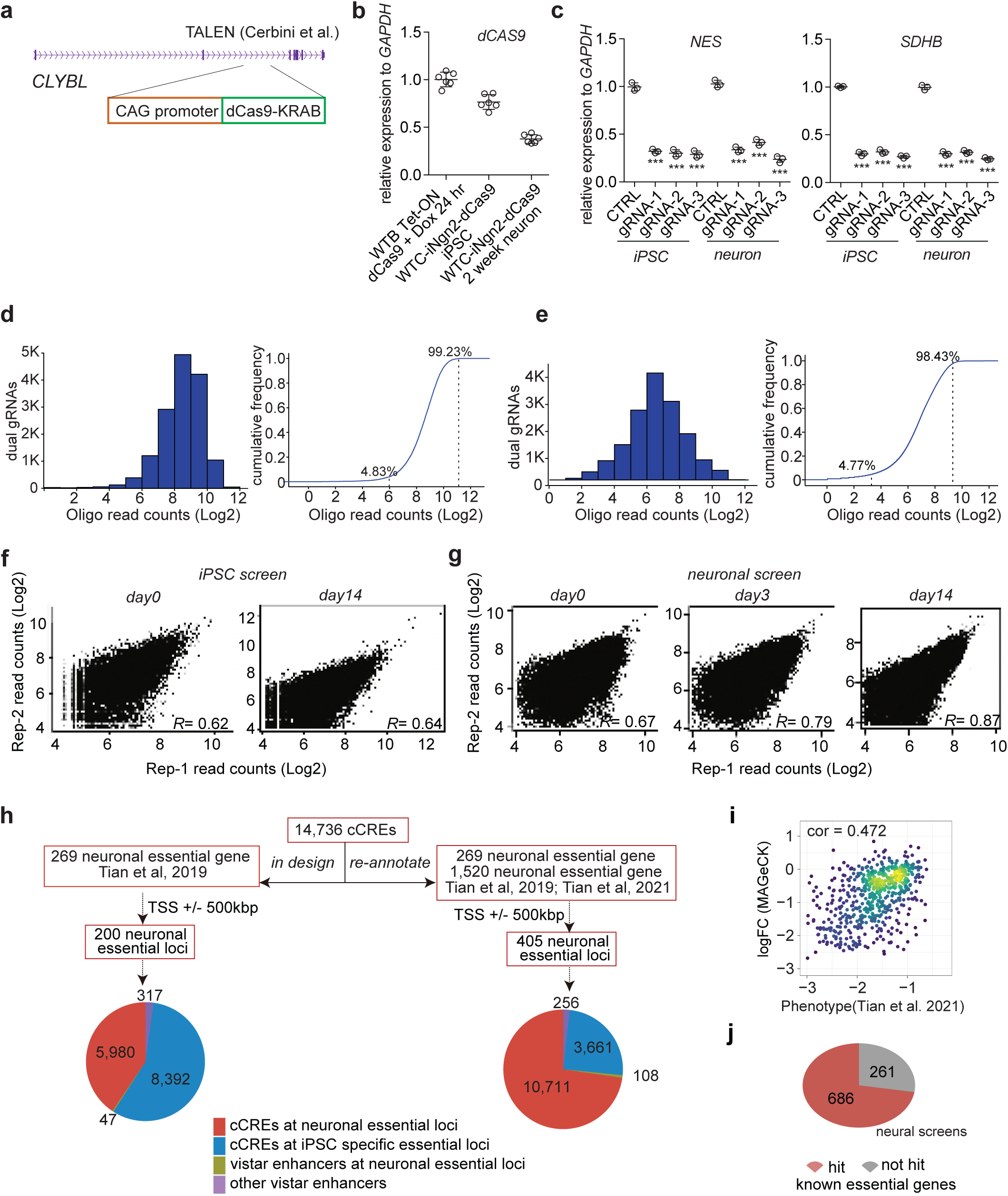
Summary of the design and results from CRISPRi screens. **a**, Schematic of genetic knock-in of dCas9-KRAB at the CLYBL safe harbor locus in i^3^N-WTC11 iPSCs. **b**, dCas9 transcription in the i^3^N-dCas9-WTC11 cell line and i^3^N-dCas9-WTC11 derived neurons. Transcription levels were compared to Tet-ON-dCas9 WTB (26971820) iPSC treated with doxycycline for 24 hrs by RT-qPCRs. **c**, CRISPRi effect targeting *SDHB* or *NES* promoter quantified by RT-qPCRs in i^3^N-dCas9-WTC11 and i^3^N-dCas9-WTC11 derived neurons. Data are shown as mean ± s.d. (n = 3, two-sided two-sample t-test, \*\*\**P* < 0.001). **d**, gRNA library distribution and recovery in the iPSC screen. **e**, gRNA library distribution and recovery in the neuronal screen. **f**, gRNA count reproducibility between biological replicates in the iPSC screen. **g**, gRNA count reproducibility between biological replicates in the neuronal screen. **h**, The chart of cCRE annotation in neuronal screen based on the 2nd study by Tian et al. **i**, Scatterplot showing the correlation of log_10_(fold change) and log_10_(phenotype score)^27^ for the neuronal essential genes. Pearson correlation score is labeled (n = 586, *P* < 2.2e-16). **j**, Count of recovered hits for neuronal essential genes identified by Tian et al.^20,27^.

**Extended Data Figure 2.**
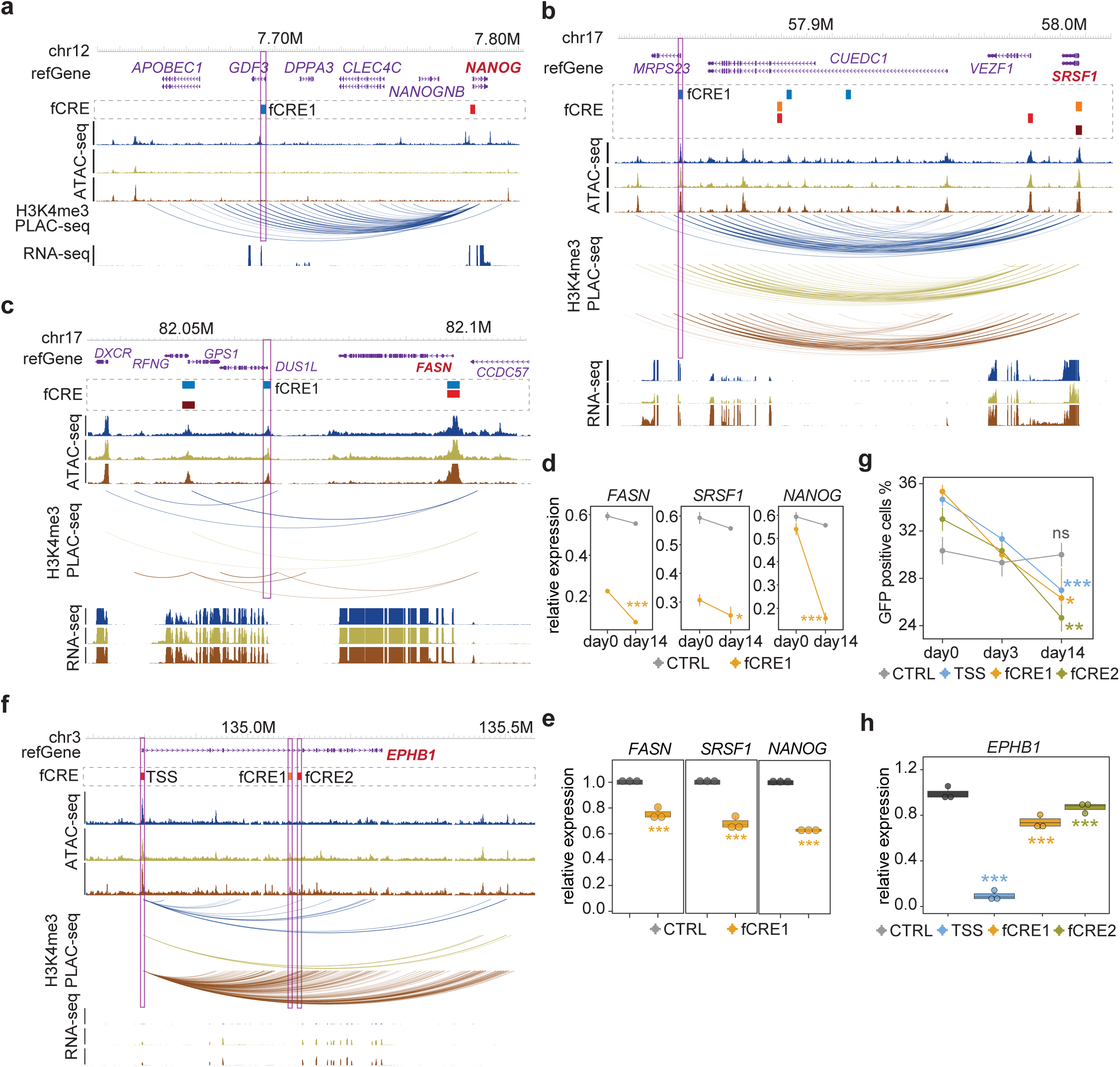
Individual CRISPRi validation for fCREs in iPSC and neuronal differentiation. **a-c**, Genome browser snapshots of the *NANOG, SRSF1* and *FASN* loci. fCREs from the iPSC, pre-differentiation, post-mitotic, and neuronal screens are colored blue, orange, red, and dark red respectively. Purple boxes indicate the validated fCREs. **d**, Survival ratio of fCRE perturbed cells by FACS in iPSC. Data are presented as mean ± s.d. (n = 3, two-tailed two sample t-test, ns: *P* > 0.05, *: *P* < 0.05, **: *P* < 0.01***: *P* < 0.001). **e.** CRISPRi effect of perturbing fCREs in iPSC quantified by RT-qPCRs. Boxplots indicate the median and interquartile range. Whiskers indicate the 5th and 95th percentiles. (n = 3, two-tailed two sample t-test). **f**, Genome browser snapshot of the *EPHB1* locus. fCREs from the iPSC, pre-differentiation, post-mitotic, and neuronal screens are colored blue, orange, red, and dark red respectively. Purple boxes indicate the validated fCREs. **g**, Survival ratio of fCRE perturbed cells by FACS in neuronal differentiation. Data are presented as mean ± s.d. (n = 3, two-tailed two sample t-test, ns: *P* > 0.05, *: *P* < 0.05, **: *P* < 0.01***: *P* < 0.001). **h.** CRISPRi effect of perturbing fCREs in neurons quantified by RT-qPCRs. Boxplots indicate the median and interquartile range. Whiskers indicate the 5th and 95th percentiles. (n = 3, two-tailed two sample t-test).

**Extended figure 3.**
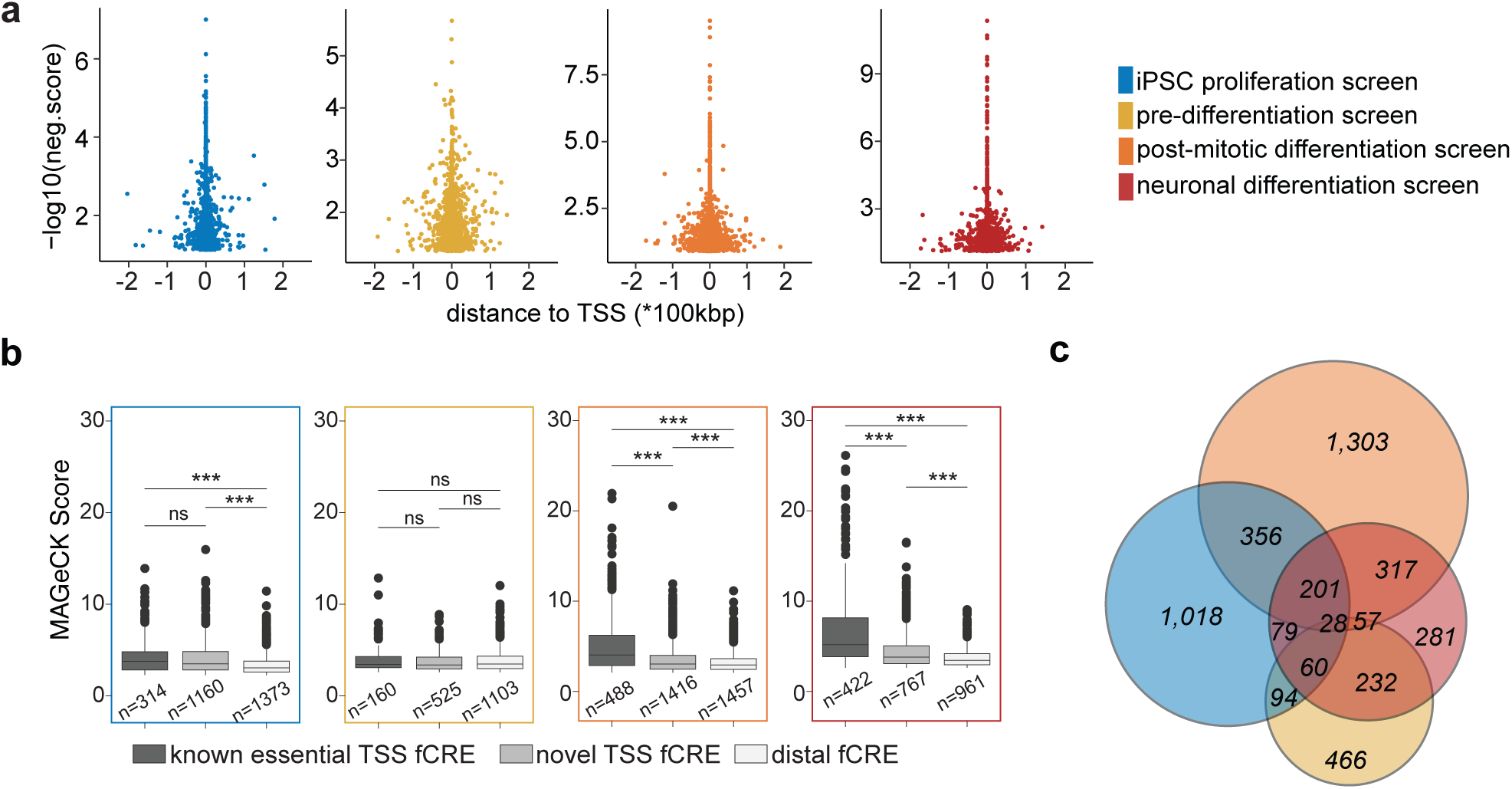
Functional annotation of fCREs in iPSC and neuronal differentiation. **a**, Scatter plot showing the MAGeCK score distribution for fCREs relative to their distance to the nearest TSS. **b**, Distribution of MAGeCK scores for known TSSs, novel TSSs, and distal fCREs for each screen (Wilcoxon test, ns: *P* > 0.05, *: *P* < 0.05, **: *P* < 0.01, ***: *P* < 0.001). **c**, Hits overlap for iPSC and neuron shared cCREs (n = 8,959) tested in the four screens.

**Extended Data Figure 4.**
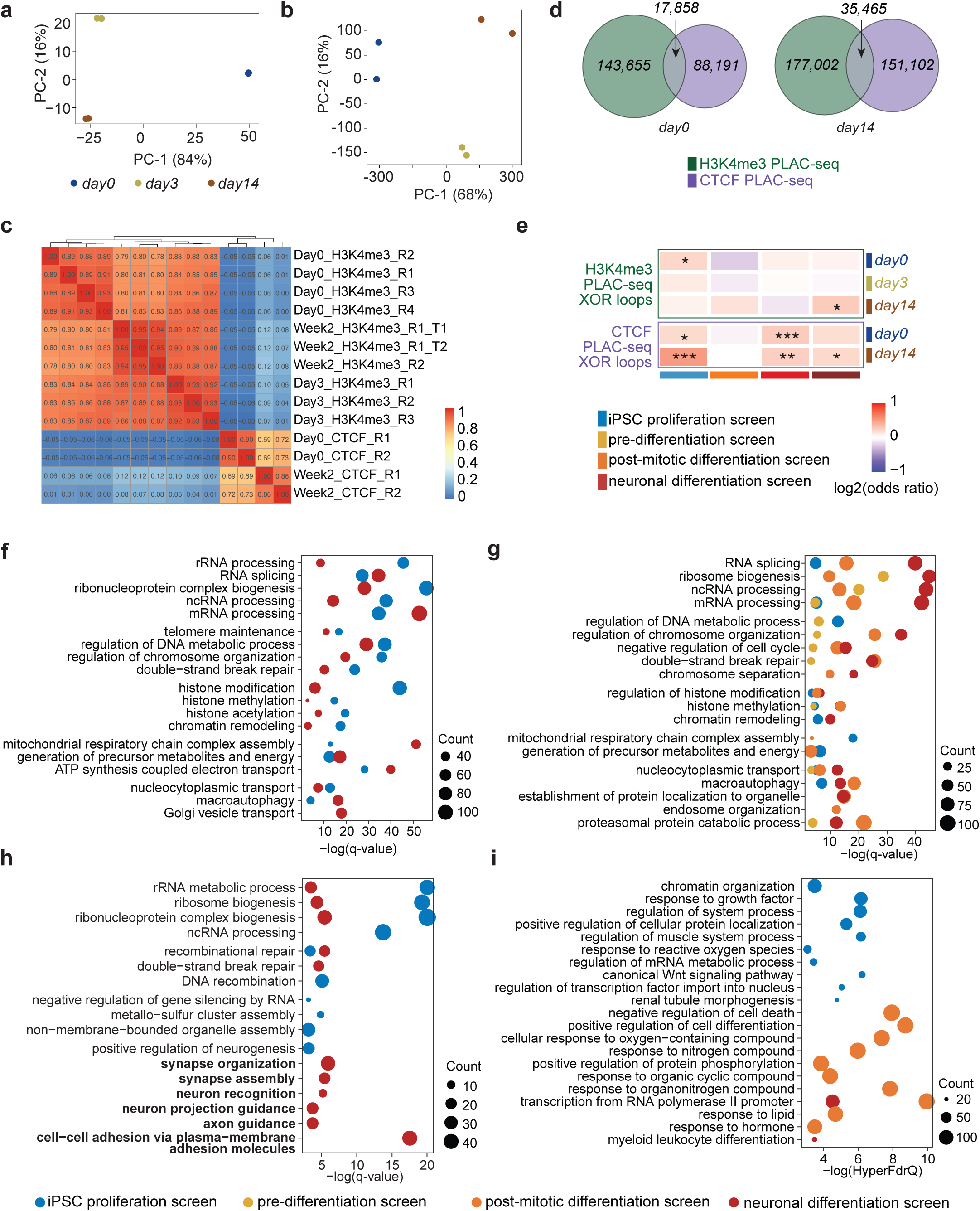
Epigenomic landscapes of neuronal differentiation and functional annotation of essential fCREs. **a**, PCA plot of RNA-seq. **b**, PCA plot of ATAC-seq. **c**, Heatmap showing the correlation of PLAC-seq datasets by HPRep. **d**, Venn diagram showing the paired-end overlapping of chromatin interactions at different time points in both H3K4me3 and CTCF PLAC-seq. **e**, Heatmap showing the association between fCREs not overlapping anchor bins and XOR chromatin interactions (N are indicated in **Supplementary Table 4b**, Fisher’s exact test, *: *P* < 0.05, **: *P* < 0.01, ***: *P*<0.001). **f**, Pathway enrichment analysis for known essential genes in library design (q-value < 0.05, iPSC, n = 1,255; neuron, n = 1,451). **g**, Pathway enrichment analysis for all TSS fCREs corresponding genes in each screen (q-value < 0.05, iPSC proliferation, n = 1,202; pre-differentiation, n = 653; post-mitotic differentiation, n = 1,794; neuronal differentiation, n = 1,172). **h**, Pathway enrichment analysis for distal fCRE target genes annotated by CTCF associated 3D chromatin interactions (q-value < 0.05, iPSC, n = 622; neuron, n = 541). **i**, Biological processes significantly enriched (HyperFdrQ < 0.05) for distal fCREs by GREAT analysis (iPSC proliferation, n = 1,251; pre-differentiation, n = 1,026; post-mitotic differentiation, n = 1,341; neuronal differentiation, n = 910).

**Extended Data Figure 5.**
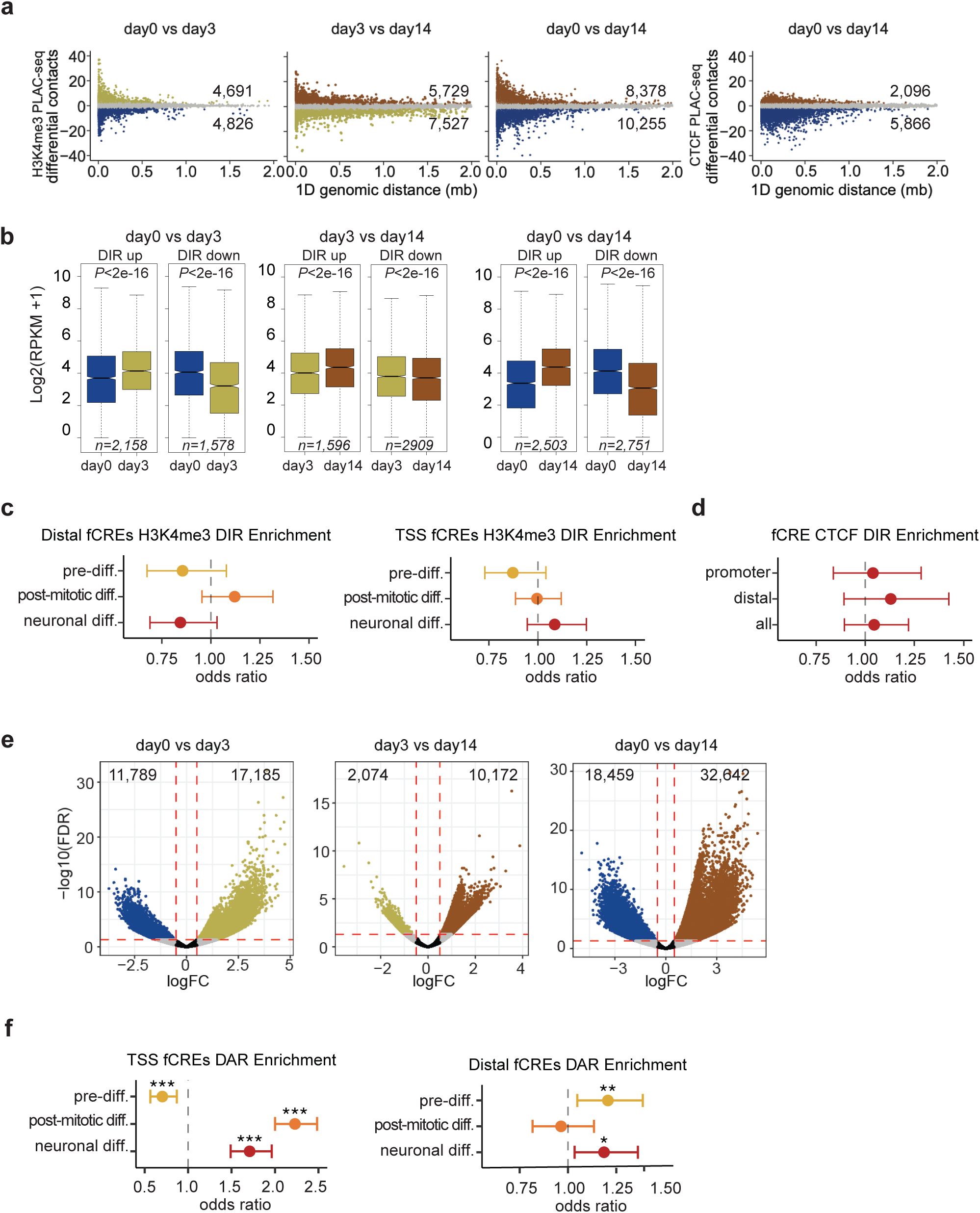
Dynamics of chromatin organization in neuronal differentiation. **a**, Scatterplot of differential chromatin interactions, where the x-axis is the 1D genomic distance (unit: Mb) between two interacting 10 Kb bins, and the y-axis is the change of contact frequency measured by +/--log_10_(*P*-value). **b**, Boxplots of gene expression (y-axis: measured by log_2_(RPKM+1)) for genes whose promoter overlaps with the differential contacts. In each box, the upper edge, horizontal center line and lower edge represent the 75th percentile, median and 25th percentile, respectively. The upper whiskers represent the 75th percentile + 1.5× the interquartile range (IQR). The lower whiskers represent the minimum values. **c-d**, Forest plot of odds ratio showing the association of distal or TSS fCREs and DIRs identified by H3K4me3 PLAC-seq (c) or CTCF PLAC-seq (d) (N are indicated in **Supplementary Table 4c**, Fisher’s exact test, *: *P* < 0.05, **: *P* < 0.01, ***: *P* < 0.001). **e**, Volcano plot of differential accessible regions for the pre-differentiation, post-mitotic, and neuronal differentiation. **f**, Forest plot of odds ratio showing the association of TSS or distal fCREs and DARs (N are indicated in **Supplementary Table 4d**, Fisher’s exact test, *: *P* < 0.05, **: *P* < 0.01, ***: *P* < 0.001).

**Extended Data Figure 6.**
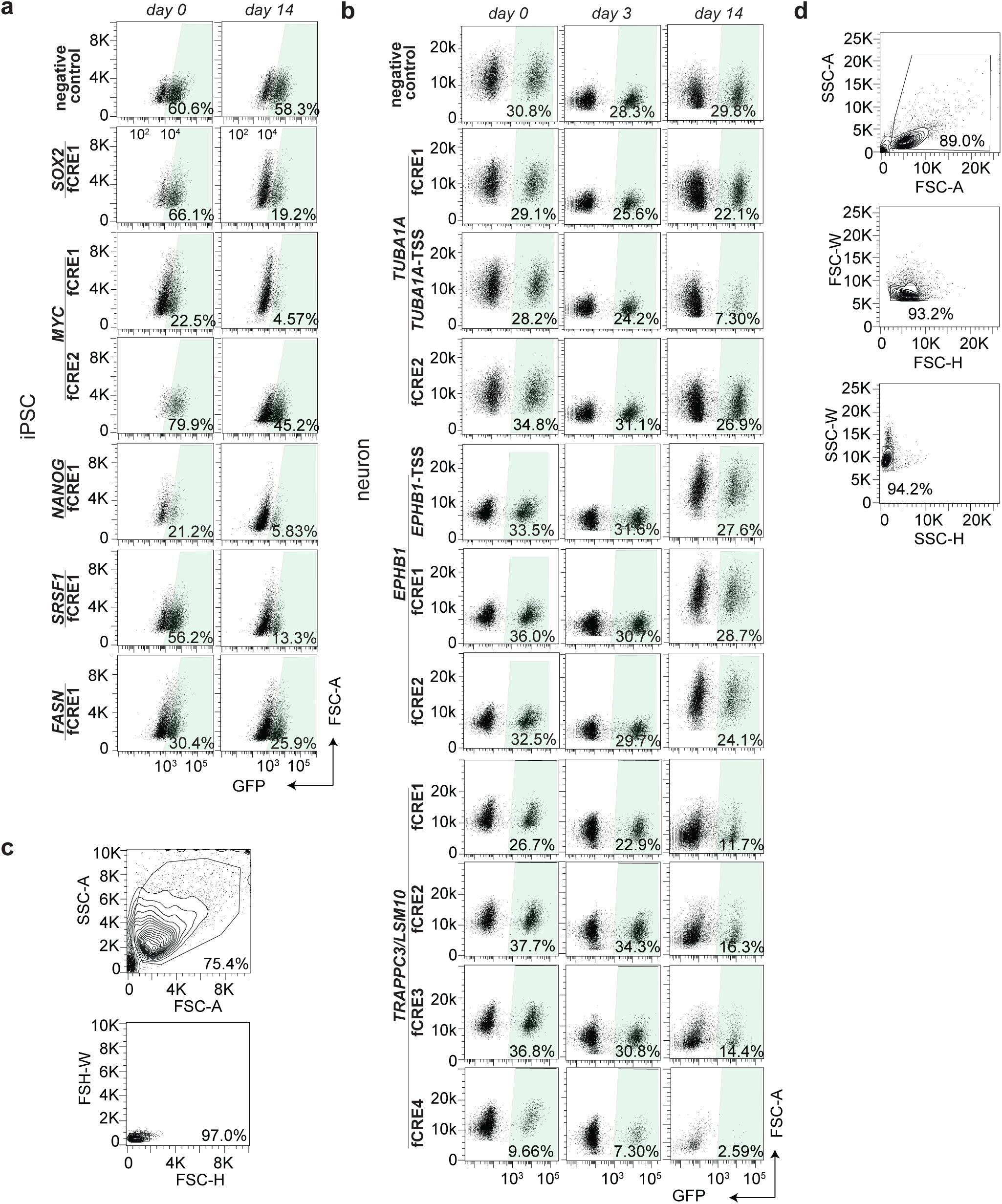
Survival ratio of fCRE perturbed cells by flow cytometry. **a**, iPSC survival after CRISPRi at the *SOX2, MYC, NANOG, SRSF1* and *FASN* fCREs. **b**, Neuronal survival ratio after CRISPRi at the *TUBA1A, EPHB1* and *LSM10* fCREs. **c-d**, Representative contour plots depicting FACS gating strategy. Cells were separated from debris of various sizes based on the forward scatter area (FSC-A) and side scatter area (SSC-A). Two singlet gates were applied using the width and height metrics of the side scatter (SSC-H versus SSC-W) and forward scatter (FSC-H versus FSC-W). All singlets are used for survival ratio analysis.

**Extended Data Figure 7.**
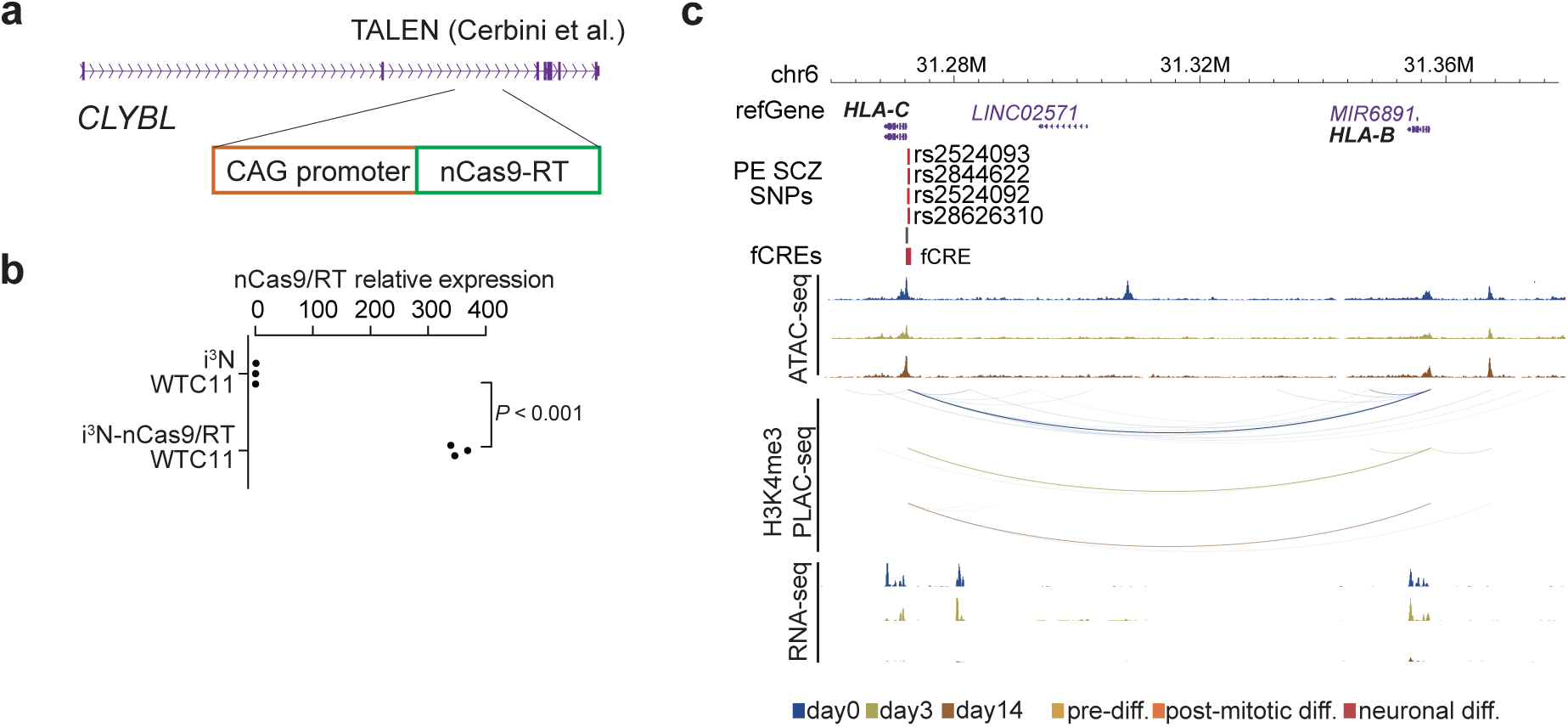
*HLA-C* association with SCZ risk. **a**, Schematic of genetic knock-in of nCas9/RT at the CLYBL safe harbor locus in i^3^N-WTC11 iPSCs. **b**, nCas9/RT expression in the i^3^N-nCas9/RT WTC11 cell line (n = 3, two-sided two sample t-test). **c**, Genome browser snapshot showing PE screened SCZ SNPs within a TSS fCRE at the *HLA-C* promoter. SNPs labeled in red indicate positive functional SNPs.

